# Distinct Vegfa isoforms control endothelial cell proliferation through PI3 kinase signalling mediated regulation of cdkn1a/p21

**DOI:** 10.1101/2020.04.01.018796

**Authors:** Martin Lange, Elvin Leonard, Nils Ohnesorge, Dennis Hoffmann, Susana F. Rocha, Rui Benedito, Arndt F. Siekmann

**Author notes:** Author for correspondence Corresponding author: Dr. Arndt F. Siekmann.

## Abstract

The formation of appropriately patterned blood vessel networks requires endothelial cell migration and proliferation. Signaling through the Vascular Endothelial Growth Factor A (VEGFA) pathway is instrumental in coordinating these processes. mRNA splicing generates short (diffusible) and long (extracellular matrix bound) Vegfa isoforms. The differences between these isoforms in controlling cellular functions are not understood. In zebrafish, *vegfaa* generates short and long isoforms, while *vegfab* only generates long isoforms. We found that mutations in *vegfaa* affected endothelial cell migration and proliferation. Surprisingly, mutations in *vegfab* specifically reduced endothelial cell proliferation. Analysis of downstream signaling revealed no change in MAPK (ERK) activation, while inhibiting PI3 kinase signaling phenocopied *vegfab* mutants. The cell cycle inhibitor *cdkn1a/p21* was upregulated in *vegfab* deficient embryos. Accordingly, reducing *cdkn1a/p21* restored endothelial cell proliferation. Together, these results suggest that extracellular matrix bound Vegfa acts through PI3K signaling to specifically control endothelial cell proliferation during angiogenesis independently of MAPK (ERK) regulation.

## INTRODUCTION

The vascular system supplies our bodies with oxygen and nutrients to ensure efficient tissue homeostasis. In order to fulfil these functions, appropriate numbers of blood vessels need to be generated in the embryo or maintained during adulthood (Adams and Alitalo, 2007; Dejana et al., 2017; Potente et al., 2011). The overproduction of blood vessels can lead to age-related macular degeneration (Mitchell et al., 2018), diabetic retinopathy (Wong et al., 2016) or cancer growth (Viallard and Larrivee, 2017). Formation of an insufficient number of blood vessels on the other hand can cause hypoxia, resulting in tissue damage as seen in atherosclerotic complication (Gimbrone and Garcia-Cardena, 2016).

Members of the vascular endothelial growth factor (VEGF) family are established regulators of vascular development (Alvarez-Aznar et al., 2017; Koch and Claesson-Welsh, 2012; Simons et al., 2016). For instance, the formation of new blood vessels from pre-exiting ones, termed angiogenesis, heavily relies on VEGFA and its receptor VEGFR2 (Kdr or Flk1). Accordingly, heterozygous *Vegfa* mutant mice die in utero with vascular defects (Carmeliet et al., 1996; Ferrara et al., 1996). Subsequent studies showed that VEGFA controls differentiation, sprouting, migration, proliferation and survival of endothelial cells (ECs). Despite the identification of several downstream players, such as Phosphoinositide 3-kinase (PI3K) (Graupera and Potente, 2013) and Mitogen-activated protein kinase (MAPK/ERK) (Simons et al., 2016), it is not known how VEGF signalling can differentially activate these pathways and how this activation might lead to the observed broad array of cellular outcomes. For example, the PI3K pathway controls EC survival in response to VEGFA in cultured cells (Gerber et al., 1998; Nakatsu et al., 2003), while *in vivo* studies in the developing mouse retina (Graupera et al., 2008) and in zebrafish embryos (Nicoli et al., 2012) suggested that PI3K signalling predominantly regulates EC migration. More recent studies, however, also suggested a function of PI3K signalling in EC proliferation during retinal development (Angulo-Urarte et al., 2018; Ola et al., 2016) and in vascular malformations (Castel et al., 2016; Castillo et al., 2019; Castillo et al., 2016). Further work in mouse and in cell culture has shown that ERK signalling downstream of VEGFA stimulates EC proliferation (Koch and Claesson-Welsh, 2012), vessel integrity (Ricard et al., 2019) and artery formation (Simons and Eichmann, 2015). Another study in zebrafish embryos, however, implicated ERK signalling mainly in regulating EC migration, being dispensable for early artery differentiation (Shin et al., 2016). Therefore, we still lack an understanding of the sequence of downstream VEGF signalling events that occur during blood vessel formation. We also do not understand how these might be triggered in different EC populations through differential ERK and PI3K signalling.

One key aspect of VEGFA biology is the existence of differentially spliced isoforms (Bowler and Oltean, 2019). Longer isoforms bind heparin and are associated with the extracellular matrix (ECM), while the short, 121 amino acid (aa) isoform is diffusible (Vempati et al., 2014). Studies in mice have shown that these isoforms differentially affect blood vessel formation. Genetically engineered mice, which only express the VEGFA165 isoform are viable (Stalmans et al., 2002), while mice expressing only VEGFA121 show angiogenesis defects (Carmeliet et al., 1999; Ruhrberg et al., 2002; Stalmans et al., 2002). Thus, the ability to associate with the extracellular matrix might change VEGFA downstream signalling and/or gradient formation, as shown in cultured cells (Chen et al., 2010).

Zebrafish contain two *vegfa* paralogs, *vegfaa* and *vegfab*, likely being generated during a genome duplication event (Taylor et al., 2003; Taylor et al., 2001). Both genes are expressed during early embryogenesis and encode differentially spliced gene products, with *vegfaa* encoding 121- and 165 aa isoforms (Gong et al., 2004; Liang et al., 1998), and *vegfab* encoding 171 and 210 aa isoforms (Bahary et al., 2007). Therefore, while *vegfaa* generates both diffusible and ECM bound isoforms, *vegfab* only generates ECM bound isoforms. Here, we have generated zebrafish mutants for *vegfaa* and *vegfab*. We find that, in agreement with studies in mice, *vegfaa* is haploinsufficient. By contrast, *vegfab* mutants showed only mild vascular defects and survived to adulthood. Surprisingly, however, EC proliferation was specifically affected in *vegfab* mutants. Using inhibitor treatments and time-lapse analysis of vascular development, we show that this phenotype was caused by impaired PI3K signalling without altering ERK activation. Further analysis of downstream targets identified the cell cycle regulator Cdkn1a/p21 as a PI3K target. Finally, we show that inhibition of PI3K signalling in the mouse retina similarly affected EC proliferation. Together, our studies support a model, in which activation of PI3K signalling downstream of ECM bound VEGFA is necessary to allow for optimal EC proliferation during angiogenesis that is independent of signalling through ERK.

## RESULTS

### Vegfaa and Vegfab mutants show distinct defects during brain blood vessel formation

In order to investigate the role of each of the zebrafish *Vegfa* homologues during angiogenesis, we generated zinc finger (for *vegfaa*) or TALEN (for *vegfab*) mutants, targeting the first exon of either gene. We recovered two frameshift mutations, leading to severely truncated proteins of 18 (*vegfaa*^*mu128*^) or 12 amino acids (*vegfab*^*mu155*^), respectively (Figure 1A, B). To assess changes in vascular morphology, we crossed both mutant lines into the *Tg(kdrl:EGFP)*^*s843*^ background, expressing EGFP in ECs (Jin et al., 2005). We first focused our analysis on the developing hindbrain vasculature (Figure 1C), which relies on VEGFR2 signaling (Bussmann et al., 2011; Fujita et al., 2011; Ulrich et al., 2011). The arterial pole of the hindbrain vasculature forms via two successive sprouting events emanating from the primordial hindbrain channels (PHBC), two laterally located veins (Bussmann et al., 2011; Fujita et al., 2011; Ulrich et al., 2011). The first sprouting event generates the medially located basilar artery (BA), while the second event subsequently generates central arteries (CtAs), which connect the PHBCs with the BA (Figure 1D-F; Video S1). Both *vegfa* genes show distinct expression domains in the brain, as shown by fluorescence in situ hybridization at 32 hpf. We detected expression of *vegfaa* dorsal to the PHBC at positions of forming CtAs (Figure S1A, arrows, B). For *vegfab*, expression was evident in the midline region, where the BA would form (Figure S1C, arrowheads, D) and dorsally of the PHBCs (Figure S1C, arrows, D).

**Figure 1.**
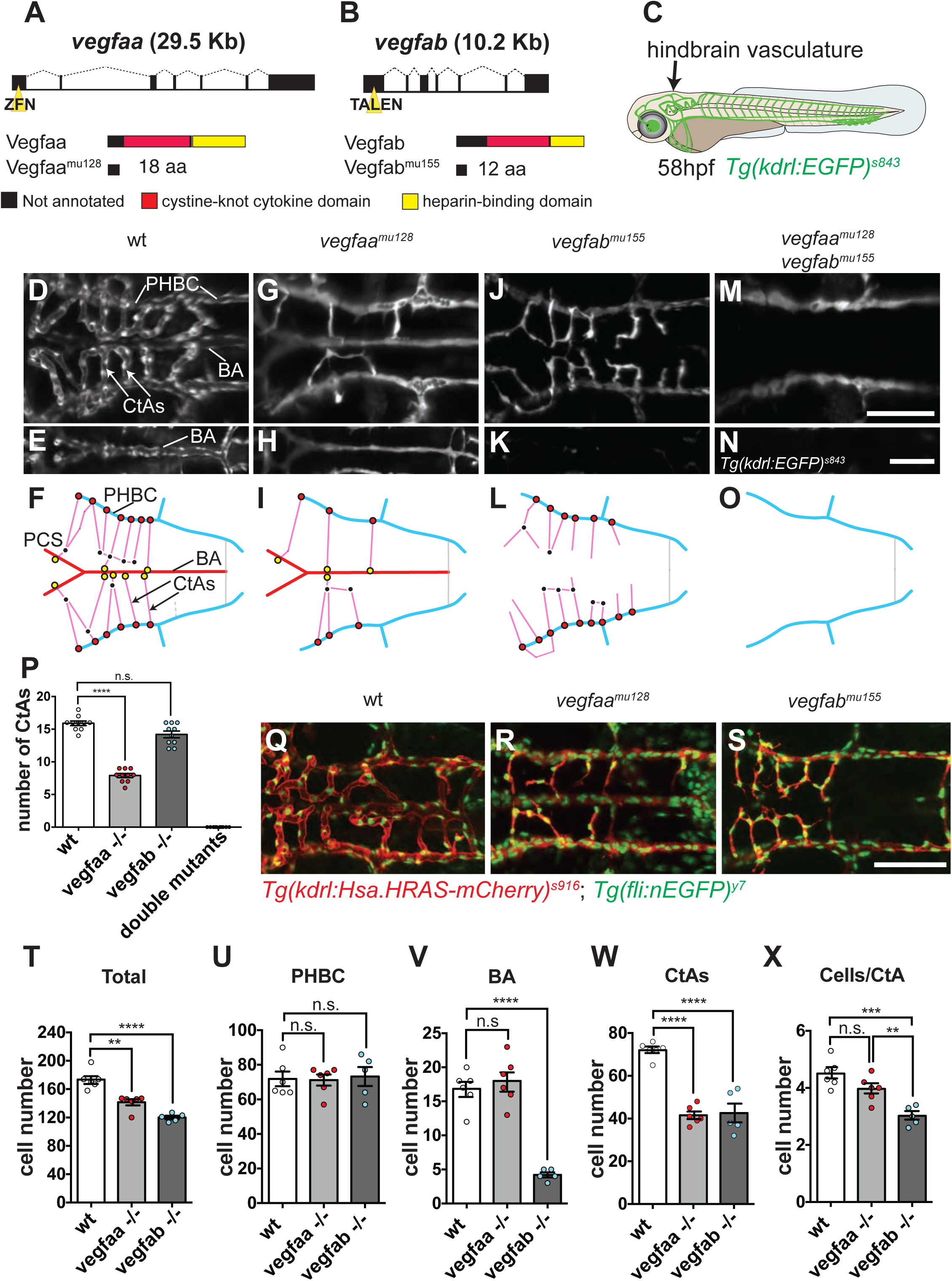
Mutations in vegfaa and vegfab affect hindbrain blood vessel formation. (A-B) Schematic presentation of zinc finger nuclease or TALEN target site at the genomic sequence for *vegfaa* (A) and *vegfab* (B). Black boxes in the gene structure represent exons and dashed lines represent introns, yellow triangle indicates the position of the targeting sites. Protein domains are displayed below in comparison with wild type protein. Black boxes represent sequences that are not annotated, red boxes are cystine-knot cytokine domains and yellow boxes are heparin binding domains. (C) Cartoon of 58 hpf embryo, arrow indicates imaged region. (D-N) Maximum intensity projection of confocal z-stacks of *Tg(kdrl:EGFP)*^*s843*^ wild type (D), *vegfaa*^*mu128*^ (G), *vegfab*^*mu155*^ (J) and double mutant (M) embryos. Smaller rectangular panels (E, H, K and N) show cropped ventral region of the maximum intensity projection for better visualization of the basilar artery (BA). Dorsal views, anterior to the left. Scale bar = 100 um. (F, I, L, O) Graphical representation of hindbrain vascular phenotypes, indicating the position of Primordial hindbrain channel (PHBC)-Central artery (CtA) connections (red filled circles), CtA-CtA connections (black dots) and CtA-BA-Posterior Communicating segments (PCS) connections (yellow filled circles). (P) Quantification of CtA number in wild type (n = 10), *vegfaa*^*mu128*^ (n = 10), *vegfab*^*mu155*^ (n = 10) and double mutant embryos (n = 10). Values are mean±s.d.; n.s = not significant; **** p<0.0001. (Q-S) Maximum intensity projections of confocal z-stacks of *Tg(kdrl:Hsa.HRAS-mCherry)*^*s916*^; *(fli1a:nEGFP)*^*y7*^ wt (Q), *vegfaa*^*mu128*^ (R) and *vegfab*^*mu155*^ embryos (S). Dorsal view, anterior to the left. Scale bar = 100 um. (T-X) Quantification of the total cell numbers (T) as well as in the PHBC (U), BA (V) CtAs (W) and cells per CtA (X), for wt embryos (n = 6) compared to *vegfaa*^*mu128*^ (n = 6) and *vegfab*^*mu155*^ (n = 5) mutants. Dots represent individual embryos; black lines indicate the mean value±s.d.; n.s = not significant; ** p>0.0022; *** p>0.0003; **** p<0.0001.

In line with these gene expression data, we observed specific vascular defects in each mutant. Homozygous *vegfaa*^*mu128*^ mutants showed a significant decrease in CtA numbers, while these were unaffected in *vegfab*^*mu155*^ mutants (Figure 1 G-L, quantified in Figure 1P; Videos S2 and S3). By contrast, the BA failed to form in these mutants, as previously reported (Rossi et al., 2016). Its formation was unaffected in *vegfaa*^*mu128*^ mutants (Figure 1G-L). To analyse whether *vegfab* can contribute to CtA formation, we analysed *vegfaa*^*mu128*^; *vegfab*^*mu155*^ double mutants. These showed a complete lack of BA and CtAs, while the PHBCs still formed (Figure 1M-O, quantified in Figure 1P). These results suggest that during brain blood vessel sprouting, *vegfaa* and *vegfab* have acquired specific regulatory elements that drive their expression in separate domains, thereby locally influencing blood vessel formation.

### Mutant hindbrain phenotypes display cell number changes in particular blood vessels

Angiogenesis requires the coordination of endothelial cell migration and proliferation (Adams and Alitalo, 2007; Hogan and Schulte-Merker, 2017; Potente et al., 2011; Schuermann et al., 2014). We therefore set out to investigate the influence of either *vegfa* gene on these processes. To do so, we quantified cell numbers using the double transgenic line *Tg(kdrl:Hsa.HRAS-mCherry)*^s916^; *Tg(fli1a:nEGFP)*^*y7*^, in which EC nuclei contain EGFP protein and membranes are labelled by virtue of mCherry expression (Hogan et al., 2009; Roman et al., 2002). After completion of BA and CtA sprouting (58 hpf time point), both mutants displayed lower total cell numbers when compared to wildtype (wt) embryos (Figure 1Q-T). Further analysis revealed that neither *vegfaa*^*mu128*^ nor *vegfab*^*mu155*^ affected PHBC cell numbers (Figure 1U). As expected, BA cell numbers were specifically affected in *vegfab*^*mu155*^ mutants (Figure 1V). The amounts of CtAs decreased about 50% in *vegfaa* mutants (Figure 1R, quantified in Figure 1P) and accordingly, total CtA cell numbers were also reduced in these mutants (Figure 1W). Surprisingly however, total CtA cell numbers were similarly decreased in *vegfab* mutants when compared to wt embryos (Figure 1W), even though *vegfab*^*mu155*^ mutants formed normal numbers of CtAs (Figure 1P). We therefore reasoned that *vegfab* might affect EC numbers in individual CtAs. Indeed, while individual CtAs in *vegfaa*^*mu128*^ mutants contained similar quantities of ECs compared to wt embryos, cell numbers per CtA in *vegfab*^*mu155*^ were significantly decreased (Figure 1X). Thus, *vegfab* specifically influenced EC numbers in blood vessel sprouts without affecting their growth. These findings suggest that the two zebrafish *vegfa* paralogs not only differentially affect blood vessel formation due to their distinct gene expression domains, but that they might also have acquired separate biological functions.

### Vegfab signalling is specifically required for EC proliferation

To test this hypothesis, we leveraged the accessibility of zebrafish for time lapse imaging. We used *Tg(kdrl:Hsa.HRAS-mCherry)* ^s916^; *Tg(fli1a:nEGFP)*^*y7*^ embryos to track the proliferative behaviours of individual ECs. In wt embryos, we observed CtA EC proliferation between 32 and 42 hpf (Figure S1E, white dots; Video S4). Ultimately, 16.5 % of CtA cells were derived from a cell division (Figure S1H, I). Although fewer CtAs sprouted in *vegfaa*^*mu128*^ mutants, the ones that formed displayed a similar percentage of CtA cells derived from proliferation when compared to wt embryos (Figure S1F, white dots, quantified in Figure S1H, I; Video S5). By contrast, *vegfab*^*mu155*^ mutants showed a reduction of cells derived from proliferation to 3.4 % (Figure S1G, white dots, quantified in Figure S1H, I; Video S6). Thus, *vegfaa* signalling cannot compensate for loss of *vegfab* signalling in stimulating EC proliferation during CtA sprouting. Together, these findings suggest that *vegfab* signalling might be critically required in controlling EC proliferation.

### VEGFab and VEGFaa differentially affect intersegmental blood vessel sprouting

To corroborate our findings, we analyzed the morphology of intersegmental blood vessels (ISVs), the first blood vessels that form via angiogenesis in zebrafish embryos (Isogai et al., 2003). In wt embryos, ISVs sprout from the dorsal aorta and anastomose in the dorsal region of the zebrafish trunk at 30 hpf (Isogai et al., 2003) (Figure S2A, B, quantified in Figure S2K). *vegfaa*^*mu128*^ heterozygous embryos showed a variable degree of stalled ISVs (Figure S2C, D, arrowhead, quantified in Figure S2K), indicating haploinsufficiency, as previously reported for VEGFA mutant mice (Carmeliet et al., 1996). ISVs failed to form in *vegfaa*^*mu128*^ homozygous embryos (Figure S2E, F, quantified in Figure S2K), as shown earlier for independently generated *vegfaa* mutants (Rossi et al., 2016). Surprisingly, neither *vegfab*^*mu155*^ heterozygous nor homozygous mutants showed defects in ISV morphology at 30 hpf (Figure S2G-J, quantified in Figure S2K). These results thus underscore a conserved role of *vegfaa* signaling during ISV formation, while *vegfab* appears to be dispensable during this process.

Motivated by our observations concerning the specific functions of *vegfaa* and *vegfab* in CtA formation, we set out to determine whether *vegfab* might similarly affect EC proliferation in ISV ECs. As previously reported, wt embryos contained either 3 or 4 cells per ISV (Figure 2A, B, quantified in Figure 2G), (Childs et al., 2002; Costa et al., 2016; Siekmann and Lawson, 2007). ISVs in heterozygous *vegfaa*^*mu128*^ mutants contained significantly fewer endothelial cells (Figure 2C, D, quantified in Figure 2G). *Vegfab*^*mu155*^ mutants displayed a reduction in ISV EC numbers (Figure 2E, F, quantified in Figure 2G) without obvious sprouting defects. Thus, similar to brain CtAs, loss of *vegfab* specifically reduces ISV cell numbers.

**Figure 2.**
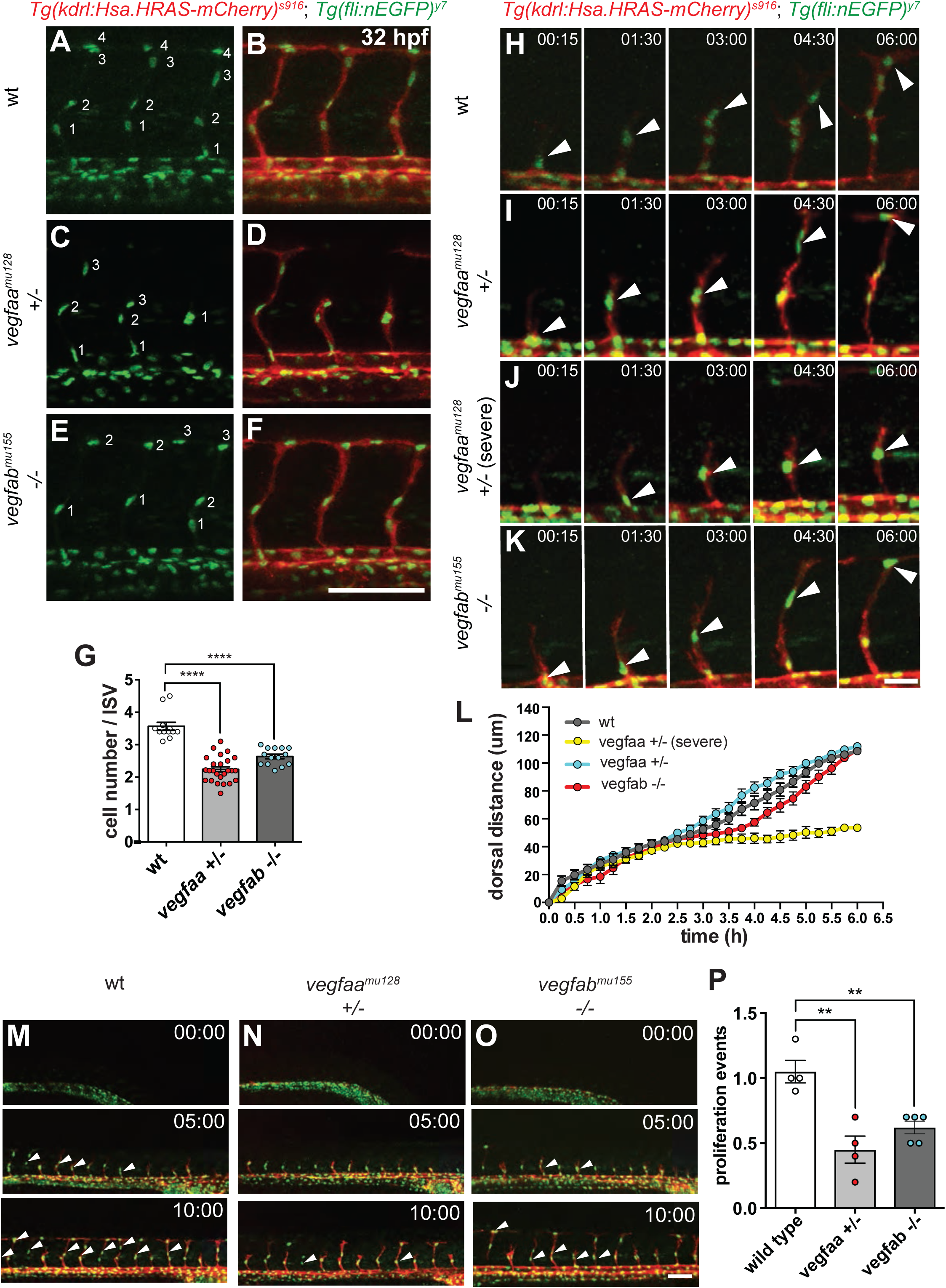
Analysis of endothelial cell migration and proliferation in the trunk vasculature of *vegfaa*^*mu128*^ and *vegfab*^*mu155*^ mutants. (A-F) Maximum intensity projections of confocal z-stacks of *Tg(kdrl:Hsa.HRAS-mCherry)*^*s916*^; *(fli1a:nEGFP)*^*y7*^ wt (A-B), heterozygous *vegfaa*^*mu128*^ (C-D) and *vegfab*^*mu155*^ (E-F) embryos; lateral view, anterior to the left. Scale bar = 100 um. (G) Quantification of endothelial cell numbers per intersegmental blood vessel sprout (ISV) in wt (n = 12), heterozygous *vegfaa*^*mu128*^ (n = 26) and *vegfab*^*mu155*^ (n = 14) mutants at 32 hpf. Dots represent individual embryos; black lines indicate the mean value ±s.d.; **** p<0.0001. (H-K) Confocal time-lapse images of individual sprouting ISV showing tip cell migration (arrowheads) in wt (H), heterozygous *vegfaa*^*mu128*^ (I), severely affected heterozygous *vegfaa*^*mu128*^ (J) and homozygous *vegfab*^*mu155*^ (K) embryos; lateral view, anterior to the left. Scale bar = 20 um. (L) Quantification of tip cell migration, measuring the dorsal movement of cell nuclei for wild type (grey; n = 13 ISVs, 4 embryos), *vegfaa*^*mu128+/-*^ (blue; n = 11 ISVs, 4 embryos), *vegfaa*^*mu128* +/-^ severe (yellow; n = 11 ISVs, 4 embryos) and *vegfab*^*mu155 -/-*^ (red; n = 11 ISVs, 4 embryos). Values are mean±s.d. (M-O) Confocal time-lapse images of 10 growing ISVs in wild type (M), *vegfaa*^*mu128+/-*^ (N) and *vegfab*^*mu155 -/-*^ (O) embryos (n=4 each) that were analyzed for cell proliferation. Arrowheads indicate cells derived from proliferation. Lateral views, anterior to the left. Scale bar = 100 um. (P) Quantification of proliferation events per ISV during 10 h of time lapse imaging from 22 - 32 hpf in wild type (n = 40 ISVs, 4 embryos), *vegfaa*^*mu128+/-*^ (n = 40 ISVs, 4 embryos) and *vegfab*^*mu155 -/-*^ (n = 50 ISVs, 5 embryos). Dots represent individual embryos; black lines indicate the mean value ±s.d.; ** p>0.0022.

We then set out to analyze how *vegfaa* and *vegfab* might affect endothelial cell migration and proliferation. To do so, we performed time-lapse imaging of Tg*(kdrl:Hsa.HRAS-mCherry)*^*s916*^; *Tg(fli1a:nEGFP)*^*y7*^ double transgenic zebrafish embryos. As determined by nuclear displacement, wt tip cells on average migrated about 100 um within 6 h (Figure 2H, arrowheads, quantified in Figure 2L; Video S7), as previously reported (Costa et al., 2016). By contrast, ISVs in *vegfaa*^*mu128*^ heterozygous fish displayed two different behaviors: Tip cells either migrated similar to wt cells (Figure 2I, quantified in Figure 2L; Video S8), or showed severe migration defects with tip cells stalling midway along the somite (Figure 2J, quantified in Figure 2L; Video S9). Therefore, lack of one *vegfaa* allele can differentially affect individual ISVs. *Vegfab* homozygous mutants showed a decrease in cell migration in the midway position, but later recovered and reached the dorsal region of the embryo at the same time as the wildtype cells did (Figure 2K, quantified in Figure 2L; Video S10). Therefore, absence of *vegfaa* signaling can have pronounced effects on EC migration, while *vegfab* mutants show only minor cell migration defects.

By contrast, EC proliferation was strongly affected in *vegfaa* and *vegfab* mutants. Time-lapse imaging revealed that in wt embryos each ISV showed around one EC proliferation event (Figure 2M, quantified in Figure 2P; Video S11). This was reduced to about half for both *vegfaa*^*mu128*^ (Figure 2N, quantified in Figure 2P; Video S12) and *vegfab*^*mu155*^ (Figure 2O, quantified in Figure 2P; Video S13) mutants. To further investigate the influence of *vegfab* on EC proliferation, we globally overexpressed Vegfab via mRNA injection. This led to an increase in EC numbers within ISVs without affecting their overall morphology (Figure S3). In summary, our findings suggest that *vegfaa* signaling drives EC migration and proliferation, while *vegfab* signaling mainly affects EC proliferation.

### Vegfab controls EC proliferation independently of ERK signaling

We reasoned that the observed differences in EC behaviors between *vegfaa*^*mu128*^ and *vegfab*^*mu155*^ mutants might enable us to dissect out signaling pathways downstream of VEGFA that specifically control proliferation. Previous studies implicated signaling through MAPK/ERK in influencing EC proliferation (Claesson-Welsh, 2016; Simons et al., 2016). We therefore set out to determine potential changes in ERK phosphorylation in *vegfab* mutants. Surprisingly, pERK antibody staining in ISVs was unchanged in *vegfab* mutant embryos (Figure 3A, B, quantified in Figure 3C). ERK signaling furthermore influences gene expression patterns within ISVs (Shin et al., 2016). We did not detect changes in either of the two reported ERK downstream genes *dll4* (Figure 3D-G) or *flt4* (Figure 3H-K) in *vegfab*^*mu155*^ mutants. Lastly, treating wt embryos with phorbol 12-myristate 13-acetate (PMA), which increased ERK phosphorylation (Figure S4), did not change EC numbers within ISVs (Figure 3L-Q, quantified in Figure 3X). PMA treatment also failed to rescue EC numbers in *vegfab*^*mu155*^ mutants (Figure 3R-W, quantified in Figure 3X). Therefore, ERK signaling does not play a major role downstream of *vegfab* signaling in controlling EC proliferation in ISVs.

**Figure 3.**
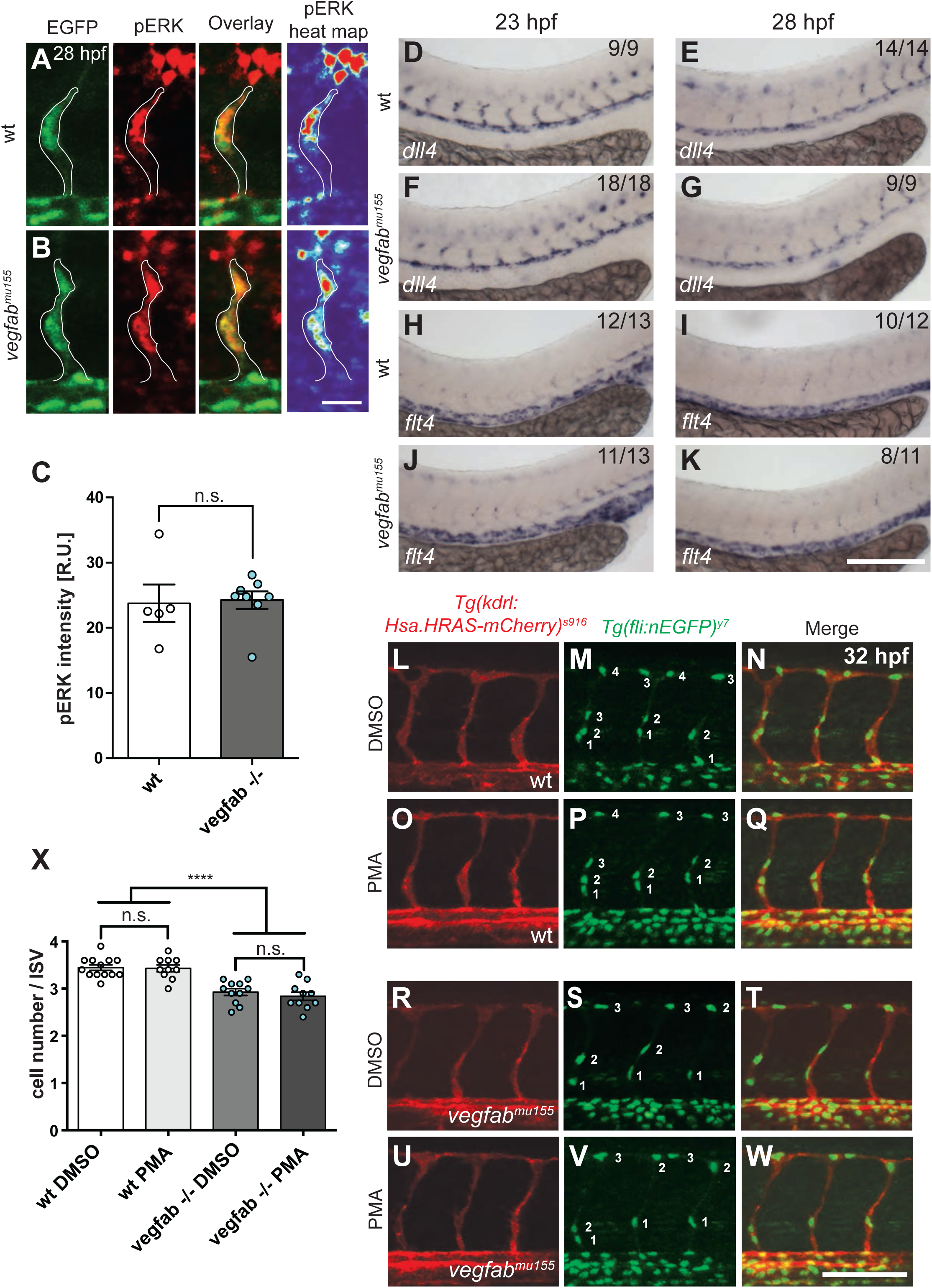
Vegfab is not essential for ERK phosphorylation or *dll4* and *flt4* expression in sprouting ISVs. (A-B) High magnification of confocal z-stack of anti-pERK antibody staining on transgenic *Tg(fli:nEGFP)*^*y7*^ embryos with quantitative heat map of pERK staining at 28 hpf for wt (A) and *vegfab*^*mu155*^ (B) embryos, lateral view, anterior to the left. Scale bar = 20 um. (C) Quantification of pERK staining intensity of every ISV cell represented in relative units (R.U.) for wt (n = 5) and v*egfab*^*mu155*^ (n = 8) at 28 hpf. Dots represent individual embryos; black lines indicate the mean value±s.d. n.s = not significant. (D-K) Whole mount in situ hybridization for dll4 (D-G) and flt4 (H-K) on wt (D-E and H-I)) and *vegfab*^*mu155*^ (F-G and J-K) embryos at 23 or 28 hpf. Lateral views, anterior to the left. Scale bar = 100 um. (L-Q) Maximum intensity projections of *Tg(kdrl:HsaHRAS-mCherry)*^*s916*^; *(fli1a:nEGFP)*^*y7*^ embryos treated with DMSO (L-N; n=13) or with 0.25 um PMA (O-Q; n=10). (R-W) Maximum intensity projections of *Tg(kdrl:HsaHRASmCherry)*^*s916*^; *(fli1a:nEGFP)*^*y7*^ v*egfab*^*mu155*^ mutant embryos treated with DMSO (R-T; n=11) or v*egfab*^*mu155*^ mutant embryos treated with PMA (U-W; n=10). Lateral views, anterior to the left. Scale bar = 100 um. (X) Quantification of endothelial cell numbers. Dots represent individual embryos; black lines indicate the mean value±s.d.; **** p<0.0001. Scale bar=100 um.

### Loss of PI3K signaling specifically affects endothelial cell proliferation

We next focused on PI3K signaling, as another important pathway downstream of VEGFA signaling (Graupera and Potente, 2013). Treating zebrafish embryos with 10 um of the PI3K inhibitor LY294002 (Vlahos et al., 1994) reduced phosphorylation of the PI3K downstream kinase AKT without affecting gross embryonic morphology (Figure S5). ISV morphology was unaffected when treating zebrafish embryos from 22 to 32 hpf (Figure 4A, D). However, similar to *vegfab*^*mu155*^ mutants, EC numbers were significantly reduced in inhibitor treated embryos when compared to DMSO treated control embryos (Figure 4B, C, E, F, quantified in Figure 4G). We then investigated whether this reduction in cell numbers was due to the reported effects of PI3K signaling on cell migration. Surprisingly, time-lapse imaging of developing ISVs did not reveal changes in cell migration upon LY294002 treatment (Figure 4H, I, quantified in Figure 4J; Videos S14 and 15). We therefore investigated EC proliferation in inhibitor treated embryos. This analysis revealed that blocking PI3K signaling led to a decrease in proliferation events (Figure 4K, L, quantified in Figure 4M; Videos S16 and S17). As LY294002 inhibits multiple PI3K isoforms in addition to other kinases (Davies et al., 2000; Workman et al., 2010), we investigated ISV formation after treating embryos with the PI3K p110 alpha isoform specific inhibitor GDC-0326 (Heffron et al., 2016). GDC-0326 treatment effectively reduced AKT phosphorylation at concentrations ranging from 10 um to 50 um (Figure S6A) and reduced ISV cell numbers without affecting ISV morphology (Figure S6B-M, quantified in Figure S6N). Thus, while not affecting ISV EC migration during short term inhibitor treatment, PI3K signaling downstream of the 110 alpha isoform plays important roles in controlling EC proliferation during developmental angiogenesis.

**Figure 4.**
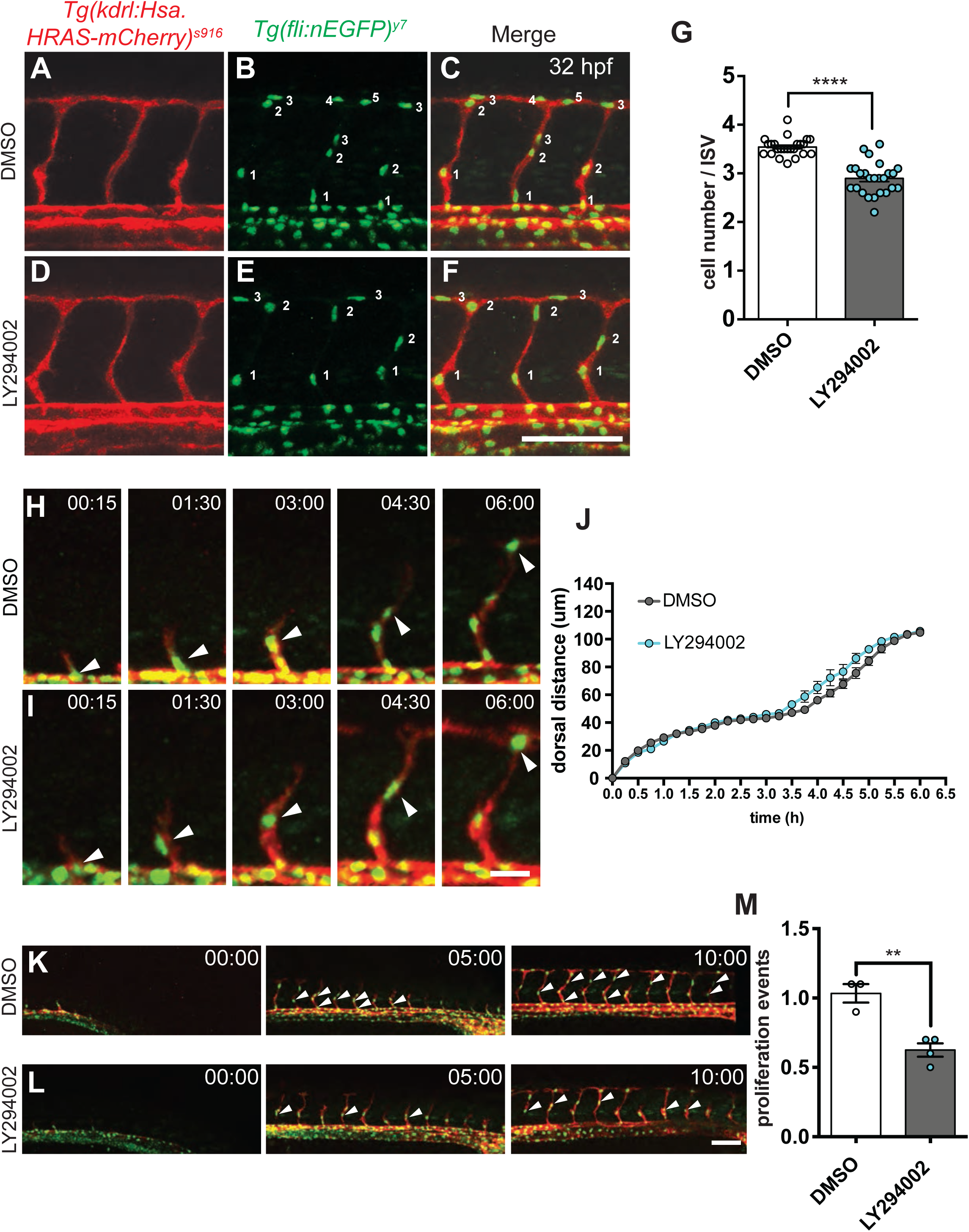
LY294002 mediated inhibition of PI3K affects endothelial cell proliferation. (A-F) Maximum intensity projection of a confocal z-stack of the trunk vasculature of double transgenic *Tg(kdrl:Hsa.HRAS-mCherry)*^*s916*^; *(fli1a:nEGFP)*^*y7*^ wt embryos treated with DMSO (A-C) or 10 uM LY294002 (D-F) at 32 hpf. Lateral views, anterior to the left. Scale bar = 100 um. (G) Quantification of the cells per ISV in DMSO treated embryos (n = 24) compared to LY294002 treated embryos(n = 25). Dots represent individual embryos; black lines indicate the mean value ±s.d. **** p<0.0001. (H, I) Confocal time-lapse images of individual sprouting ISV showing tip cell migration (arrowheads) in wt embryos treated with DMSO (H) or LY294002 (I) starting from 22 hpf for 6 h. Scale bar = 20 um. (J) Quantification of tip cell migration, measuring the dorsal movement of cell nuclei for wild type embryos treated with DMSO (grey; 11 ISVs,3 embryos) or LY294002 (blue; 11 ISVs, 4 embryos); Values are mean±s.d. (K, L) Confocal time-lapse images of 10 growing ISVs in embryos treated with DMSO (K) or with LY294002 (L) that were analyzed for cell proliferation. Arrowheads indicate cells derived from proliferation. Scale bar = 100 um. (M) Quantification of proliferation events in ISVs during 10 h of time lapse imaging from 22 - 32 hpf after DMSO (n = 30 ISVs, 3 embryos) or LY294002 (n = 40 ISVs, 4 embryos) treatment. Dots represent individual embryos; black lines indicate the mean value±s.d.; ** p>0.0022.

### Cdkn1a/p21 is a downstream target of vegfab/PI3K signaling in regulating EC proliferation

To identify *vegfab* target genes that might lead to altered proliferative behaviors, we used Fluorescence-activated cell sorting (FACS) to sort GFP and mCherry positive ISV and arterial ECs from *vegfab* morpholino injected embryos using a triple transgenic strategy (Figure 5A). We then analyzed expression of genes implicated in cell cycle regulation. This analysis revealed mRNA upregulation of the negative cell cycle regulator *cdkn1a/p21* in ECs of *vegfab* morpholino injected embryos (Figure 5B). We furthermore determined the influence of PI3K inhibition on Cdkn1a/p21 protein expression in cultured human umbilical vein endothelial cells (HUVEC). Overall Cdkn1a/p21 protein levels were unchanged after LY294002 treatment (Figure 5C). However, we detected a striking change in Cdkn1a/p21 protein localization from cytoplasmic towards nuclear (Figure 5D-I, quantified in Figure 5J). Nuclear localization has been implicated in mediating the cell growth inhibiting function of Cdkn1a/p21 (Zhou et al., 2001). Thus, both *vegfab* and PI3K signaling can influence the abundance and localization of Cdkn1a/p21 in ECs.

**Figure 5.**
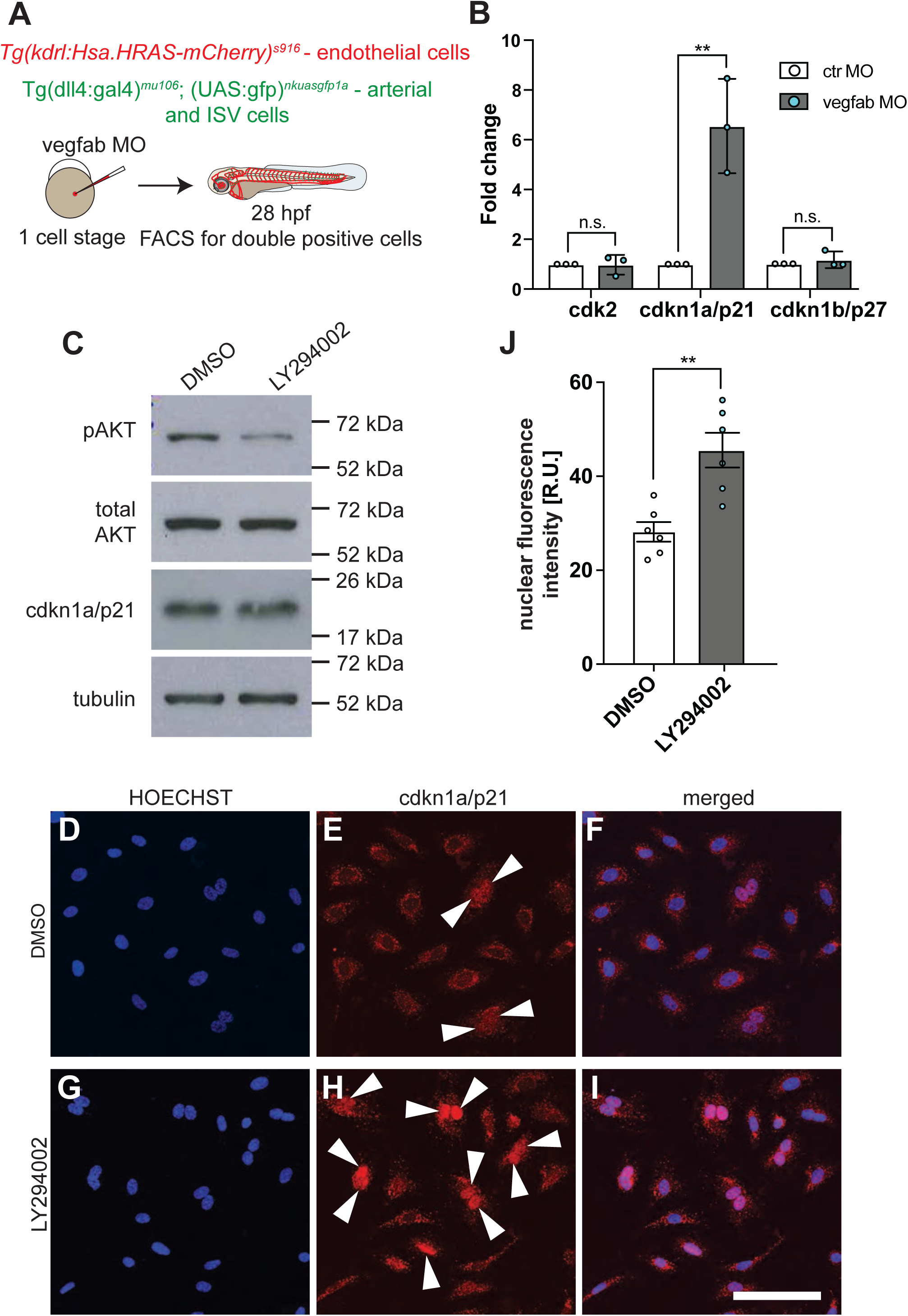
Vegfab / PI3K signaling influences Cdkn1a/p21 expression and localization. (A) Cartoon depicting experimental set up for FACS and qPCR experiment. Triple transgenic embryos *Tg(kdrl:Hsa.HRAS-mCherry)*^*s916*^; *(dll4:gal4)*^*mu106*^; *(UAS:gfp)*^*nkuasgfp1a*^ were injected with vegfab or control morpholino at the 1 cell stage and raised to 28 hpf. Embryos were dissociated and GFP/mCherry positive cells were FACS sorted. (B) Expression of cell cycle regulators cdk2, cdkn1a/p21 and cdkn1b/p27 comparing cells from vegfab morpholino injected embryos with control morpholino injected embryos, which were set as 1 (n = 3 independent experiments). Expression data are shown as fold change; dots represent individual experiments; black lines indicate the mean value±s.d.; ** p>0.0022. (C) Western Blot of proteins from Human Umbilical Vein Endothelial Cells showing decrease in AKT phosphorylation, while no change in Cdkn1a/p21 protein amount following LY294002 treatment was detected. Representative blot of n=3 is shown. (D-I) Confocal images of Human Umbilical Vein Endothelial Cells treated with DMSO (D-F) or LY294002 (G-I). HOECHST staining labels nuclei, Cdkn1a/p21 protein in red, scale bar is 200 um. (J) Quantification of nuclear fluorescence of Cdkn1a/p21 protein in DMSO (n=6 experiments) or LY294002 (n=6 experiments) treated Human Umbilical Vein Endothelial Cells in relative units (R.U.). Dots represent individual experiments; black lines indicate the mean value±s.d.; ** p>0.0022.

Based on these findings, we investigated whether knocking down Cdkn1a/p21 could rescue the proliferation defects observed upon loss of *vegfab* or PI3K signaling. Injection of Cdkn1a/p21 morpholino into wt embryos caused a mild increase in ISV EC numbers (Figure 6A, B, quantified in Figure 6E). Injecting Cdkn1a/p21 morpholinos into *vegfab*^*mu155*^ mutants led to a rescue of EC numbers back to wt levels (Figure 6C, D, quantified in Figure 6E). Similarly, injection of Cdkn1a/p21 morpholino prior to PI3K inhibitor treatment increased ISV EC numbers back to wt levels (Figure 6F-I, quantified in Figure 6J). Therefore, reducing Cdkn1a/p21 levels can rescue the proliferation defects observed upon loss of *vegfab* and PI3K signaling.

**Figure 6.**
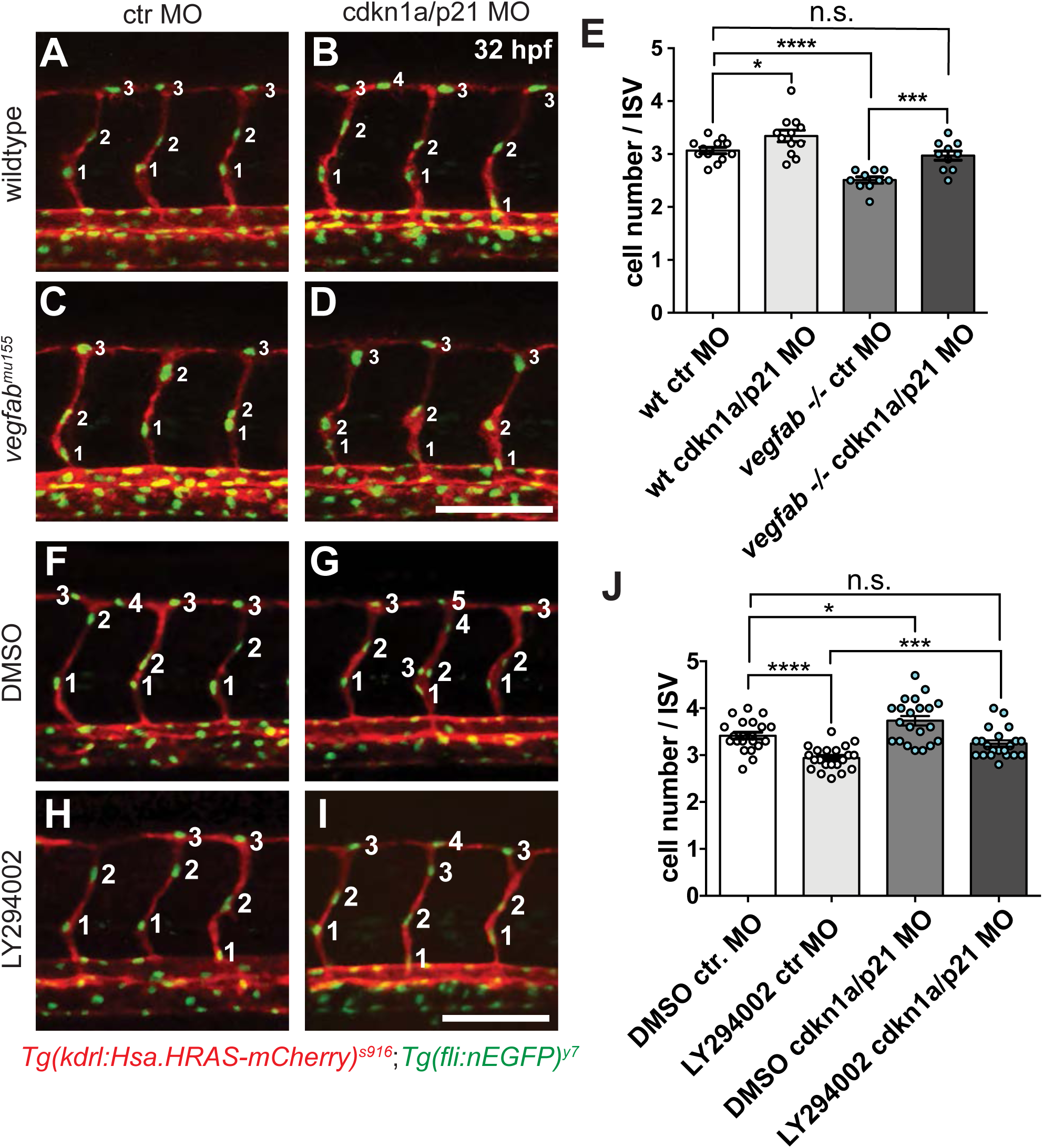
Cdkn1a/p21 knock down in zebrafish embryos rescues proliferation defects resulting from *vegfab* deficiency or PI3K inhibition. (A-D) Maximum intensity projection of a confocal z-stack of the trunk vasculature of double transgenic *Tg(kdrl:Hsa.HRAS-mCherry)*^*s916*^; *Tg(fli1a:nEGFP)*^*y7*^ wild type embryos (A-B) or *vegfab*^*mu155*^ embryos (C-D) at 32 hpf, injected with either control morpholino (A, C) or Cdkn1a/p21 morpholino (B, D). Lateral view; anterior to the left. Scale bars = 100 um. (E) Quantification of cells per ISV in wt (n=12) and *vegfab*^*mu155*^ (n=10) embryos injected with control or Cdkn1a/p21 morpholino (MO). Dots represent individual embryos; black lines indicate the mean value ±s.d. n.s = not significant; * p>0.05; *** p>0.0003; **** p<0.0001. (F-I) Maximum intensity projection of a confocal z-stack of the trunk vasculature of 32 hpf double transgenic *Tg(kdrl:Hsa.HRAS-mCherry)*^s916^; *Tg(fli1a:nEGFP)*^*y7*^ embryos injected with ctr. morpholino (F, H) or cdkn1a/p21 morpholino (G, I) that were also treated with either DMSO (F-G) or 10 uM LY294002 (H-I). Lateral view; anterior to the left. Scale bars = 100 um. (J) Quantification of cells per ISV at 32 hpf for Ctr. MO + DMSO, Ctr. morpholino + LY294002, cdkn1a/p21 MO + DMSO and cdkn1a/p21 MO + LY294002 (n = 21 embryos in each group); black lines indicate the mean value ±s.d. n.s = not significant; * p>0.05; *** p>0.0003; **** p<0.0001.

### Inhibition of PI3 kinase signaling during retinal angiogenesis reduces endothelial cell proliferation

To investigate whether PI3K signaling influences EC proliferation in other angiogenic settings, we analyzed blood vessel development in the mouse retina. Here, new blood vessels sprout from the optic nerve towards the periphery of the retina after birth. Intraperitoneal injection of the PI3k 110 alpha subunit-specific inhibitor GDC-0941 effectively reduced AKT phosphorylation in ECs after 24 h of treatment, as analyzed in lung lysates from P6 mice (Figure S7A). It furthermore reduced the phosphorylation of S6 kinase, a downstream target of PI3K signaling (Chung et al., 1994) (Figure S7B-E). Incorporation of EdU was strongly reduced in inhibitor treated embryos, both at the angiogenic front (Figure 7A-D, quantified in G) and in the vein region of the retina (Figure 7E, F, quantified in H), where ECs continue to proliferate. Thus, similar to our observations in zebrafish embryos, PI3K signaling affects endothelial cell proliferation during mouse retinal angiogenesis.

**Figure 7.**
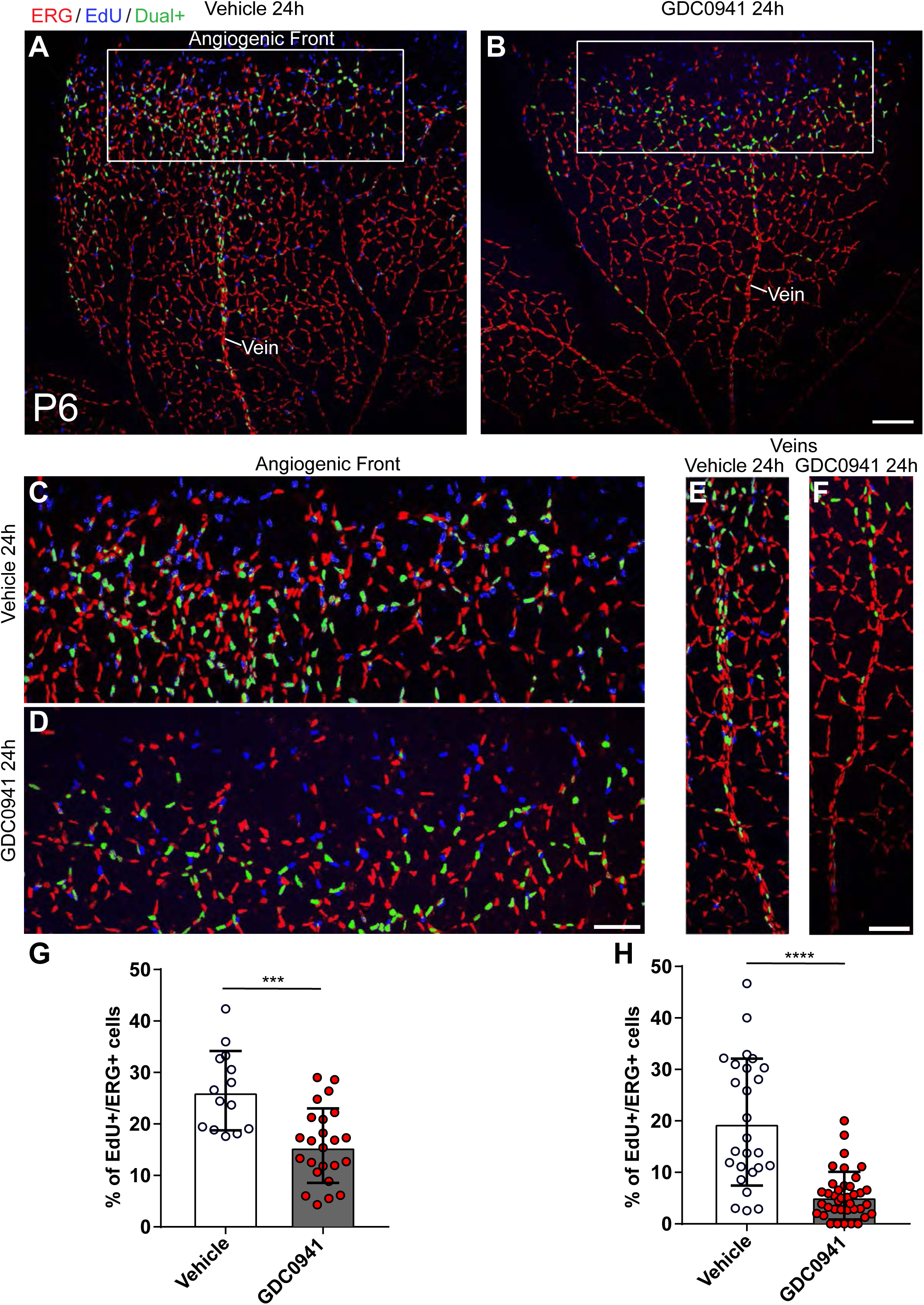
Inhibiting PI3 kinase signaling during retinal angiogenesis reduces endothelial cell proliferation. (A) Representative confocal micrograph of a vehicle treated P6 retina. Endothelial cell nuclei are labelled in red by ERG, EdU incorporation in blue and overlay of red and blue channels is pseudocolored in green. Boxed area is magnified in (C). (B) Representative confocal micrograph of a GDC-0941 treated P6 retina. Boxed area is magnified in (D). Scale bar is 150 um. (C) Magnified image of angiogenic front in vehicle treated retina. (D) Magnified image of GDC-0941 treated retina. (E) Magnified image of vehicle treated vein. (F) Magnified image of GDC-0941 treated vein. (G) Quantification of percentage of EdU positive ERG positive cells in vehicle or GDC-0941 treated retinas at the angiogenic front. Each dot represents a retina flank. For vehicle treated group n=4 (7 retinas), for GDC-0941 treated group n=5 (10 retinas). (H) Quantification of percentage of EdU positive ERG positive cells in vehicle or GDC-0941 treated retinas in the vein area. Each dot represents a retina flank. For vehicle treated group n=4 (7 retinas), for GDC-0941 treated group n=5 (10 retinas).

## DISCUSSION

Tissue vascularization requires coordinated cellular responses to growth factors, such as Vegfa, in order to generate appropriate amounts of new ECs, allow them to sprout into avascular areas and ultimately differentiate into the correct numbers of arteries, capillaries and veins. How a single ligand can control the multitude of cellular responses necessary to achieve these tasks is an outstanding question in the field. Our work on the duplicated Vegfa ligands in zebrafish identifies a specific function of PI3 kinase signaling in response to ECM bound *vegfab* ligands in regulating EC proliferation. This was an unexpected discovery, since signaling through MAPK/ERK was thought to be the main driver of EC proliferation downstream of Vegfa signaling (Koch and Claesson-Welsh, 2012; Meadows et al., 2001; Simons et al., 2016; Srinivasan et al., 2009; Takahashi et al., 1999). What might be the reason for this discrepancy? So far, animal models carrying mutations in components of the Vegf pathway showed simultaneous defects in EC proliferation and migration. For instance, zebrafish mutants in *kdrl, plcg* or treated with the Vegf pathway inhibitor SU5416 display an absence of ERK phosphorylation (Fish et al., 2017; Shin et al., 2016), together with a reduction in ISV outgrowth and cell numbers. Similarly, mutants in *vegfaa* show a severe reduction in EC numbers and migration during ISV formation (Jin et al., 2017), phenotypes also seen in *VEGFA* mutant mice (Carmeliet et al., 1996; Ferrara et al., 1996). Therefore, it has been difficult in animal models to dissect out the individual contribution of a given gene or its downstream pathway components to either process.

Specific manipulations of ERK signaling in cultured ECs showed a requirement for DNA synthesis downstream of VEGFR2 activation, a process that was independent of PI3K activity (Takahashi et al., 1999). However, deletion of ERK1 in mice did not result in vascular phenotypes (Pages et al., 1999), while knocking out ERK2 specifically in ECs in an ERK1^-/-^ background affected EC migration and proliferation (Srinivasan et al., 2009). Our study together with the study of Shin et al. (Shin et al., 2016) suggests that in developing ISVs, ERK signaling rather controls EC migration instead of proliferation. A similar specific influence of ERK signaling on migration was reported in a HUVEC tube formation assay and for tumor ECs (Mavria et al., 2006). These results suggest that ERK phosphorylation can influence EC migration and/or proliferation depending on the developmental setting and the EC type analyzed.

Our work points towards a critical role of PI3K signaling for EC proliferation during ISV outgrowth. We find that blocking PI3K using different inhibitors selectively prevented ISV EC proliferation without influencing cell migration. This is in contrast to previous studies showing that PI3K signaling specifically affected EC migration in developing mouse embryos and in cultured ECs, with only minor effects on EC proliferation (Graupera et al., 2008; Takahashi et al., 1999). A previous study in zebrafish ISVs also showed that blocking PI3K signaling using the LY294002 inhibitor affected cell migration without reducing EC proliferation (Nicoli et al., 2012). We directly imaged dividing cells using time-lapse imaging, while Nicoli et al. determined differences in BrdU incorporation. It might therefore be that PI3K signaling is important for cytokinesis with less effects on DNA synthesis, as previously shown in *Dictyostelium discoideum* cells (Janetopoulos et al., 2005). In addition, differences in the duration of PI3K inhibition will affect EC behaviors. We blocked PI3K signaling for 10 hours, while Graupera et al. analyzed vascular phenotypes after several days of removing PI3K 110alpha kinase function from ECs (Graupera et al., 2008).

Other studies have shown that PI3K signaling downstream of VEGF receptor signaling can lead to an increase in EC proliferation. Dayanir et al. generated a chimeric VEGF receptor 2 that could be activated using CSF-1 (Dayanir et al., 2001). When PI3K signaling was compromised, CSF-1 stimulation of this receptor failed to induce EC proliferation. Another study showed that Y1212 in VEGFR2 was important to control PI3K signaling pathway activation upstream of myc-dependent EC proliferation (Testini et al., 2019). In line with our observations in the mouse retina, blocking PI3K signaling in this setting reduced the number of phospho-histone 3 positive ECs (Ola et al., 2016) and led to a reduction in EdU incorporation (Angulo-Urarte et al., 2018). Importantly, activating mutations in PIK3CA can lead to venous malformations that are characterized by increased EC proliferation (Castel et al., 2016; Castillo et al., 2019; Castillo et al., 2016). Together, these studies suggest a more important role of PI3K signaling in controlling EC proliferation downstream of Vegf signaling than previously anticipated. However, further work will be necessary to precisely determine in which EC populations and at which stages of the cell cycle PI3K signaling is required. Of note, we still observed EC proliferation in PI3K inhibitor treated or *vegfab* mutant embryos, suggesting the existence of other signaling pathways contributing to ISV EC proliferation.

Why are Vegfab ligands specifically affecting PI3K signaling and EC proliferation without a major influence on EC migration? One reason for this might due to the existence of differentially spliced Vegfa isoforms. In other species, all Vegfa isoforms are being generated by the same gene, while in zebrafish only *vegfaa* can generate both short and long, ECM binding, isoforms (Bahary et al., 2007). *Vegfab* exclusively generates ECM binding isoforms. This setting allowed us to determine the unique effects of ECM binding Vegfa isoforms on EC behaviors during embryogenesis. Previous studies in developing mice (Ruhrberg et al., 2002; Stalmans et al., 2002), disease settings (Brash et al., 2019; Cheng et al., 1997; Guo et al., 2001; Kazemi et al., 2016) and in cultured ECs (Chen et al., 2010; Delcombel et al., 2013; Fearnley et al., 2016; Herve et al., 2005; Park et al., 1993; Shiying et al., 2017) carefully investigated the effects of the different VEGF isoforms on cellular behaviors (for review see (Peach et al., 2018; Woolard et al., 2009)). Some studies suggested that there are no differences in the ability of various VEGFA isoforms to support EC proliferation (Ruhrberg et al., 2002), while others showed that ECs cultured on ECM derived from cells expressing VEGF189 or VEGF206 proliferated more strongly than those cultured in the presence of VEGF165 (Park et al., 1993). Our studies support the latter findings by showing that matrix bound Vegfab isoforms stimulate EC proliferation. We furthermore find that EC proliferation in *vegfab*^*mu155*^ mutants is more strongly affected in actively sprouting ECs in the forming CtAs and ISVs. We hypothesize that this might be due to the fact that during angiogenic growth ECs degrade the ECM, possibly releasing bound VEGF molecules, which would make them available to the invading ECs. Thus, the ability of long VEGF isoforms to associate with the ECM could provide a readily available growth factor pool within tissues.

Cell cycle regulation is a key prerequisite for proper embryonic development and tissue homeostasis and is regulated by a stoichiometric competition between proliferation promoting cyclinD expression that is induced by mitogens and inhibitory cdkn1a/p21 (Pack et al., 2019). Previous work showed that cdkn1a/p21 levels vary between cells in a given population and that these levels can determine the proportion of cells that actively cycle (Overton et al., 2014). Furthermore, increased levels of cdkn1a/p21 after mitosis can lead to cellular quiescence, while cells with lower cdkn1a/p21 levels continue to proliferate (Spencer et al., 2013). A similar balance of cdkn1a/p21 might control the number of proliferating ECs during ISV sprouting. Indeed, when comparing different ISVs, we observe a variation in proliferating ECs. In cdkn1a/p21 knockdown embryos, a greater number of ECs proliferate, while a reduction in *vegfab* signaling increases expression of cdkn1a/p21 and causes fewer ECs to proliferate. Recently, increases in ERK signaling were shown to upregulate cdkn1a/p21 expression in ECs in the mouse retina, resulting in cell cycle arrest (Pontes-Quero et al., 2019). Our results suggest that PI3K signaling downstream of *vegfab* similarly controls the amount and/or localization of cdkn1a/p21, thereby determining the amount of actively cycling ECs during angiogenesis.

Of note, cdkn1a/p21 can also influence cell migration via the inhibition of Rho/ROCK (Rho-associated kinase) signaling (Kreis et al., 2019; Tanaka et al., 2002). During tumorigenesis, upregulation of cdkn1a/p21 can induce growth arrest, allowing breast cancer cells to become more migratory and invasive (Qian et al., 2013). Previous studies showed that activation of the PI3K target Akt results in cdkn1a/p21 cytoplasmic localization, releasing cdkn1a/p21’s growth inhibiting function (Zhou et al., 2001). Our results show a similar effect of PI3K signaling on cdkn1a/p21localization in ECs. Therefore, proper activation of PI3K signaling upstream of cdkn1a/p21 might be important for controlling the switch between EC migration and proliferation. A better understanding of the mechanisms controlling this switch will further our understanding of tissue morphogenesis and help to manipulate aberrant blood vessel development in beneficial ways.

## MATERIALS AND METHODS

### Zebrafish strains

Zebrafish were maintained as described previously (Westerfield, 1993). Transgenic lines and mutants used were *Tg(kdrl:EGFP)*^*s843*^, *Tg(kdrl:Hsa.HRAS-mCherry)*^*s916*^, *Tg(fli1a:nEGFP)*^*y7*^, *Tg(dll4:gal4)*^*mu106*^, *Tg(UAS:GFP)*^*nkuasgfp1a*^, *vegfaa*^*mu128*^ (this study), *vegfab*^*mu155*^ (this study). References for zebrafish transgenic lines can be obtained on zfin.org. Adult zebrafish used in this study to generate embryos were between 1-2 years of age. Embryos were not selected for gender. All animal experiments were performed in compliance with the relevant laws and institutional guidelines and were approved by local animal ethics committees of the Landesamt für Natur, Umwelt und Verbraucherschutz Nordrhein-Westfalen.

### Morpholino injections and drug treatment

Morpholinos were obtained from Gene-Tools and dissolved in distilled water. Morpholinos (MOs) targeting *cdkn1a/p21* (3 ng/embryo) (Sidi et al., 2008), *vegfab* (5 ng/ embryo) (Bahary et al., 2007) and a standard control MO (Hasan et al., 2017) have been described previously. To inhibit PI3K, embryos were dechorionated and incubated in 10 uM LY-294002 hydrochloride (SIGMA) for 10 hours prior to imaging. To specifically inhibit PI3K 110 alpha, embryos were dechorionated and incubated in 10 uM, 25 uM or 50 uM of GDC-0326 (Cayman Chemical). To activate ERK signalling, embryos were dechorionated and incubated in 0.25 uM Phorbol-12-myristat-13-acetate (PMA) (SIGMA) for 4 hours prior to imaging.

### Live imaging, confocal microscopy and image processing

For *in vivo* imaging, live embryos were mounted in 1 % or 1.5 % low-melting-point agarose in E3 embryo medium with 168 mg l^−1^ tricaine (1x) for anaesthesia and 0.003 % phenylthiourea to inhibit pigmentation. Imaging was carried out on an inverted Leica SP5, SP8 or a Zeiss LSM780 confocal microscope using a 20× dry objective. A heated microscope chamber at 28.5 °C was used for recording time-lapse videos. Stacks were taken every 15-20 min with a step size of 2 um. Confocal stacks and time-lapse videos were analysed using IMARIS Software (Bitplane).

### Fluorescence-activated cell sorting (FACS) of arterial ECs from zebrafish embryos

Arterial ECs were obtained from triple-transgenic zebrafish embryos, in which *Tg(kdrl:Hsa.HRAS-mCherry)*^*s916*^ exhibits pan-endothelial expression, and the combination of *Tg(dll4:gal4)*^*mu106*^; *(UAS:GFP)*^*nkuasgfp1a*^ labels arterial ECs within this population. Embryos were deyolked in calcium free Ringer’s solution and treated with 0.5% Trypsin-EDTA (Gibco, 15400-054) containing 50 mg/ml collagenase Type IV (Gibco,17104-019) and dissociated by constant pipetting. The reaction was stopped by adding 5% FBS and the cells were pelleted by centrifugation. The pelleted cells were washed with 1x HBSS buffer (Gibco, 14185052) and passed through 40-micron nylon filters. FACS was performed on the cell suspension for arterial (mCherry+/GFP+) and the venous (mCherry+) cells at room temperature. Sorted cells were collected in RLT buffer (Qiagen) for RNA isolation.

### Quantitative Polymerase Chain Reaction (qPCR)

RNA was isolated with RNeasy Plus Micro kit (Qiagen) from whole zebrafish embryos or sorted cells and reverse transcribed with iScript cDNA Synthesis Kit (BioRad). For qPCR, cDNA produced from 15 ng RNA and 8 pmol of each forward and reverse primer per reaction and Power SYBR Green PCR Master Mix (Applied Biosystems) were used. Relative expression was quantified by ΔΔCt method using RPL13A as an endogenous control. The expression in control sample (unsorted cells/ctr MO injected embryos) was set as 1. ΔCt values were used for statistical analysis. The following primers were used:

Cdk2-fwd: 5’ -CGGAGGGCACTGTTTCCTGGAG- 3’

Cdk2-rev: 5’ -ACATTTGCCCAAGAAGGTCTCTGCC- 3’

Cdkn1a/p21-fwd: 5’ -AGCTTCAGGTGTTCCTCAGCTCCT- 3’

Cdkn1a/p21-rev: 5’ -CCGGCCCGAAAAGACTCCGC- 3’

Cdkn1b/p27-fwd: 5’ -TACAGCCGATCGGAGCGGGG- 3’

Cdkn1b/p27-rev: 5’ -TGACACGATGAGTCGAGACAGGAGC- 3’

RPL13A-fwd: 5’ - TCTGGAGGACTGTAAGAGGTATGC

RPL13A-rev: 5’ - AGACGCACAATCTTGAGAAGCAG

### Generation of *vegfaa*^*mu128*^ and *vegfab*^*mu155*^ mutant zebrafish

Zinc-finger nucleases (ZFNs) against *vegfaa* were designed as previously described (Siekmann et al., 2009). In the *vegfaa*^*mu128*^ allele, 7 nucleotides were deleted in exon 1 at the ZFN target site, resulting in an early stop codon after 18 amino acids. TALEN mutagenesis targeting *vegfab* was performed as described previously (Sugden et al., 2017). In the *vegfab*^*mu155*^ allele, 1 nucleotide was deleted and 2 nucleotides were inserted in exon 1 at the TALEN target site. This resulted in an early stop codon after 12 amino acids.

### Genotyping

Primers for genotyping *vegfaa* were:

Vegfaa-fwd: 5’ -GCTTTCTTAATTGTTTTGAGAGCCAG- 3’

Vegfaa-rev: 5’ –GGTGTGGGCTATTGCATTTC- 3’

PCR products were digested with BccI (NEB). Fragment sizes are 116 bp + 123 bp for wild type allele and 239 bp for *vegfaa*^*mu128*^ allele.

Primers for genotyping vegfab were:

Vegfab-fwd: 5’ -GGACCAACATGGGATTCTTG- 3’

Vegfab-rev: 5’ -GGGTGGTCAGATATGCTCGT- 3’

PCR products were digested with BsrI (NEB). Fragment sizes are 188 bp + 221 bp for wild type allele and 409 bp for *vegfab*^*mu155*^ allele.

### Cloning of *vegfab171* and overexpression studies

*Vegfab171* was amplified with primers VEGFabattB1 ggggacaagtttgtacaaaaaagcaggctGTTAAAAACGGGCAACGGCGG and VEGFab attB2 ggggaccactttgtacaagaaagctgggtTCACCTCCTTGGTTTGTCACATCTGC from zebrafish 24 hpf cDNA. cDNA was generated using RNA that was isolated with RNeasy Plus Micro kit (Qiagen) from whole zebrafish embryos and reverse transcribed with iScript cDNA Synthesis Kit (BioRad). BP reaction was performed according to the manufacturer’s instructions (ThermoFisher). Clones were verified by sequencing. *Vegfab* was then transferred into pCSDest (Villefranc et al., 2007) using LR cloning (ThermoFisher). Plasmid DNA was digested using NotI (NEB) and mRNA was generated using mMessage machine in vitro transcription kit (ThermoFisher). 50 pg of mRNA was injected into 1-cell stage zebrafish embryos.

### In situ hybridization and fluorescence in situ hybridizations (FISH) with antibody staining

Whole-mount in situ hybridization was performed as previously described (Thisse and Thisse, 2008). Whole mount FISH combined with EGFP antibody staining was performed in *Tg(kdrl:EGFP)*^*s843*^ line as described previously (Kochhan and Siekmann, 2013). Previously described probes were for *vegfaa* (Lawson et al., 2002), *vegfab* (Bahary et al., 2007), *dll4* (Siekmann and Lawson, 2007), *flt4* (Lawson et al., 2001).

### Western blotting of zebrafish proteins

Dechorionated zebrafish embryos were de-yolked in Ginzburg buffer (55 mM NaCl, 1.8 mM KCl, 1.25 mM NaHCO_3_) and lysed in Laemli buffer (20 embryos in 80 ul). Either 10 or 20 ul of sample were separated by 12 % SDS-PAGE and transferred onto PVDF membrane (Millipore). After blocking with 5 % Milk powder (Roth) in TBST, membranes were incubated with anti-phospho-p44/42 MAPK (Thr202/Tyr204) (1:1000; #8544; Cell Signaling), anti-p44/42 MAPK antibody (1:2000; #9102, Cell Signaling) and anti-αActin antibody (1:5000; A-5060, Sigma), Phospho-Akt (Ser473) (1:2000; #4060; Cell Signaling), Akt (1:1000; #9272; Cell Signaling). Primary antibodies were detected using mouse IgG HRP linked whole Ab (1:4000; NXA931; GE healthcare).

### Immunostaining of zebrafish embryos

Immunostaining for phospho-p44/42 MAPK was performed as described previously (Costa et al., 2016).

### Mouse retina

To inhibit PI3K signaling, Pictilisib (GDC-0941, Selleckchem) stock solution was prepared by dissolving 8.5 mg of powder in 63 ul DMSO. Before injection, 10 ul of the stock solution was diluted in 190 ul of corn oil to get a final concentration of 6,8 ug/ul. 25 ul of this solution (or vehicle only) was injected IP into each pup at P5 (55 mg/kg) and again 16 h later, before collecting the tissues at P6 (injections at -24 h and -8 h time points). To detect proliferating cells actively synthesizing DNA, EdU (Invitrogen - A10044) was injected IP 4 h before sacrifice; the signal was developed with the Click-it EdU Alexa Fluor 647 Imaging Kit.

### Immunohistochemistry of mouse retinae

For mouse retina immunostaining, eyes were collected at the indicated time points and fixed in 4% PFA in PBS for 1h at room temperature (RT). After two PBS washes, retinas were micro-dissected and stained as described previously (Pontes-Quero et al., 2019). Briefly, retinas were blocked and permeabilized with 0.3% Triton X-100, 3% FBS and 3% donkey serum in PBS. Samples were then washed twice in PBLEC buffer (1 mM CaCl_2_, 1 mM MgCl_2_, 1 mM MnCl_2_ and 1% Triton X-100 in PBS). Biotinylated isolectinB4 (Vector Labs, B-1205, diluted 1:50) or primary antibodies (see below) were diluted in PBLEC buffer and tissues were incubated in this solution for 2 h at RT or overnight at 4°C. After five washes in blocking solution diluted 1:2, samples were incubated for 1 h at RT with Alexa-conjugated secondary antibodies (Molecular Probes). After two washes in PBS, retinas were mounted with Fluoromount-G (SouthernBiotech). To detect EdU-labelled DNA, an additional step was performed before mounting using the Click-It EdU kit (Thermo Fisher, C10340). Primary antibodies were used against the following proteins: Erg (AF-647, Abcam ab196149, 1:100), Phospho-S6 Ribosomal Protein (Ser235/236) (Cell Signalling Ab #4856, 1:100). The following secondary antibodies were used: Donkey anti-rabbit Cy3 (1:400, 711-167-003, Jackson Immunoresearch) and Streptavidin Alexa 405 (1:400, S-32351, Thermofisher).

### Western blot analysis of mouse proteins

For the analysis of protein expression, dissected organs were transferred to a reagent tube and frozen in liquid nitrogen. On the day of the immunoblotting the tissue was lysed with lysis buffer [(Tris-HCl pH=8 20 mM, EDTA 1 mM, DTT 1 mM, Triton X-100 1% and NaCl 150 mM, containing protease inhibitors (P-8340 Sigma) and phosphatase inhibitors (Calbiochem 524629) and orthovanadate-Na 1 mM)] and homogenized with a cylindrical glass pestle. Tissue/ cell debris was removed by centrifugation, and the supernatant was diluted in loading buffer and analyzed by SDS–PAGE and immunoblotting. Membranes were blocked with BSA and incubated with primary antibodies diluted 1/1000 against Cdh5/VE-cadherin (BD Biosciences 555289), Phospho-Akt (Cell Signalling, #4060S), Akt (Cell Signaling #9272S) or β-Actin (Santa Cruz Biotechnologies, sc-47778).

### Microscopy of mouse retina

We used a Leica TCS SP8 confocal with a 405 nm laser and a white laser that allows excitation at any wavelength from 470nm to 670nm. All images shown are representative of the results obtained for each group and experiment. Littermates were dissected and processed under exactly the same conditions. Comparisons of phenotypes or signal intensity were made with pictures obtained using the same laser excitation and confocal scanner detection settings. Images were processed using ImageJ/Fiji and Adobe Photoshop.

### Quantitative analysis of retinal vasculature

Single low magnification (10x lens) confocal fields of immunostained retinas were quantified with Fiji/ImageJ. Each microscopy field contained hundreds of ECs, and the relative or absolute number of cells in each field is indicated in the charts by a dot. As indicated in figure legends, microscopy images from several animals and retinas were used for the phenotypic comparisons and quantifications. Vascular IsolectinB4+ area and Erg+ or Edu+ cells were quantified semiautomatically using custom Fiji macros. Endothelial cell density (EC number/mm^2^) was measured as the number of Erg+ cells relative to the vascularized IsolectinB4+ area in each field. The frequencies of Erg+ cells (ECs) in S-phase (EdU+) was determined as the ratio of double-positive cells to the total number of Erg+ cells per field.

### Statistical analysis

Two groups of samples with a Gaussian distribution were compared by unpaired two-tailed Student *t*-test. Comparisons among more than two groups were made by ANOVA followed by the Turkey pairwise comparison. Column statistics were performed on data sets to check for normal distribution and appropriate tests to determine significance were performed using the Prism7 software. Each experiment was performed at least three times. Graphs represent mean +/- SD as indicated, and differences were considered significant at p < 0.05. All calculations were done in Excel and final datapoints analyzed and represented with GraphPad Prism. The sample size was chosen according to the observed statistical variation and published protocols. The experiments were not randomized, investigators were not blinded to allocation during experiments and outcome assessment and sample sizes were not predetermined.

## MATERIALS AVAILABILITY

All reagents and zebrafish lines generated in this study are available from the Lead Contact with a completed Materials Transfer Agreement.

## ACKNOWLEDGEMENTS

We would like to thank Reinhild Bussmann, Mona Finch Stephen, Nadine Greer and Bill Vought for excellent fish care. In addition, we would like to thank Roman Tsaryk and Zeenat Diwan for critically reading of the manuscript. We are grateful to Federica Lunella for help with the mouse retina dissection and immunohistochemistry. This work was funded by the Max-Planck-Society (A.F.S.), the Deutsche Forschungsgemeinschaft (DFG SI-1374/4-1, DFG SI-1374/5-1 and DFG SI-1374/6-1; A.F.S.) and start-up funds from the Cardiovascular Institute of the University of Pennsylvania Perelman School of Medicine (A.F.S.). We further acknowledge support from the Department of Cell and Developmental Biology of the University of Pennsylvania (A.F.S.). Work in R.B.’s lab was funded by the Ministerio de Economía, Industría y Competitividad (MEIC: SAF2017-89299-P and RYC-2013-13209) and the European Research Council (ERC-2014-StG – 638028 AngioGenesHD).

## AUTHOR CONTRIBUTIONS

M.L. and A.F.S. conceived the experiments and analyzed the data. A.F.S. supervised the work. M.L. analyzed *vegfaa* and *vegfab* mutants, performed drug treatmens, pERK stainings and p21 knockdown experiments. M.L. also performed all cell culture experiments. A.F.S. cloned *vegfab* and performed overexpression experiments. N.O. generated the *vegfaa* and *vegfab* mutants. E.L. performed FACS of zebrafish endothelial cells and qPCR experiments. D.H. analyzed data. S.F.R. performed mouse retina experiments and analyzed data. R.B. analyzed data of mouse retina. All authors discussed experiments and commented on the manuscript.

## FIGURE LEGENDS

**Supplementary Figure S1.**
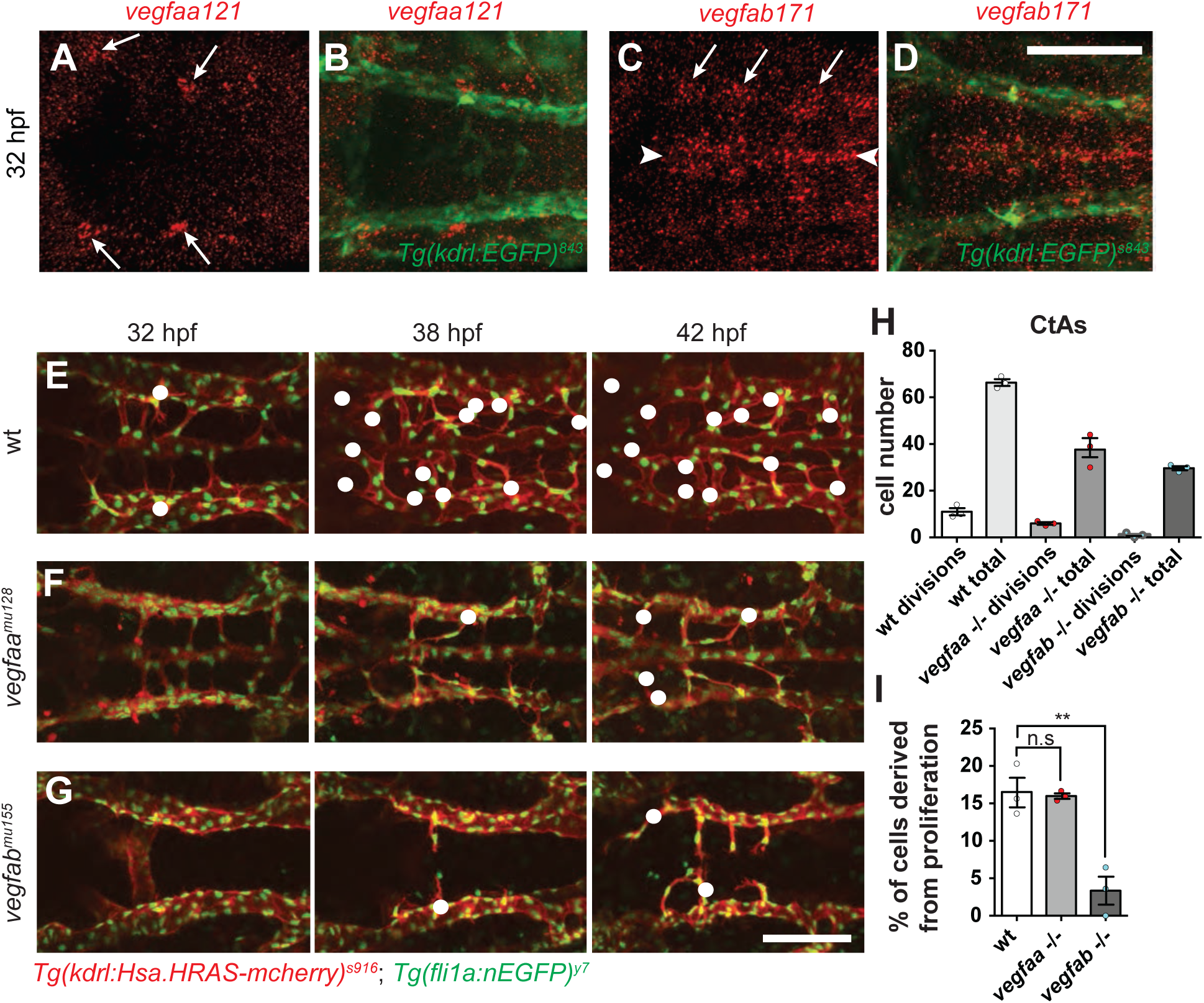
Distinct sprouting and proliferation defects in *vegfaa* or *vegfab* mutants correlate with *vegfaa* and *vegfab* expression domains. (A-D) Fluorescent in situ hybridization combined with an anti-EGFP antibody staining showing the distribution of vegfaa121 (red, A-B), vegfab171 (red, C-D) and EGFP driven by *Tg(Kdrl:EGFP)*^*s843*^ (green, B, D) at 32 hpf. Dorsal views anterior to the left. Arrows indicate the expression domains of vegfaa and vegfab. Scale bar = 100 um. (E-G) Still images of confocal time-lapse of zebrafish hindbrain development in live *Tg(kdrl:Hsa.HRAS-mCherry)*^*s916*^; *Tg(fli1a:nEGFP)*^*y7*^ embryos between 32 and 42 hpf, shown for wild type (E), vegfaa^mu128^ (F) and vegfab^mu155^ (G). Dorsal view, anterior to the left, Scale bar = 100 um. White dots mark cells within CtAs derived from proliferation. (H) Quantification of cell numbers derived from proliferation and total cell numbers in wild type, *vegfaa*^*mu128*^ and *vegfab*^*mu155*^ embryos (n = 3 embryos for each group). Dots represent individual embryos; black lines indicate the mean value±s.d. n.s. = not significant; * p>0.05; ** p>0.0022; **** p<0.0001. (I) Quantification of cells derived from proliferation normalized to the total cell number comparing wild type, *vegfaa*^*mu128*^ and *vegfab*^*mu155*^ (n = 3 embryos for each group). Dots represent individual embryos; black lines indicate the mean value±s.d. n.s = not significant; ** p>0.0022.

**Supplementary Figure S2.**
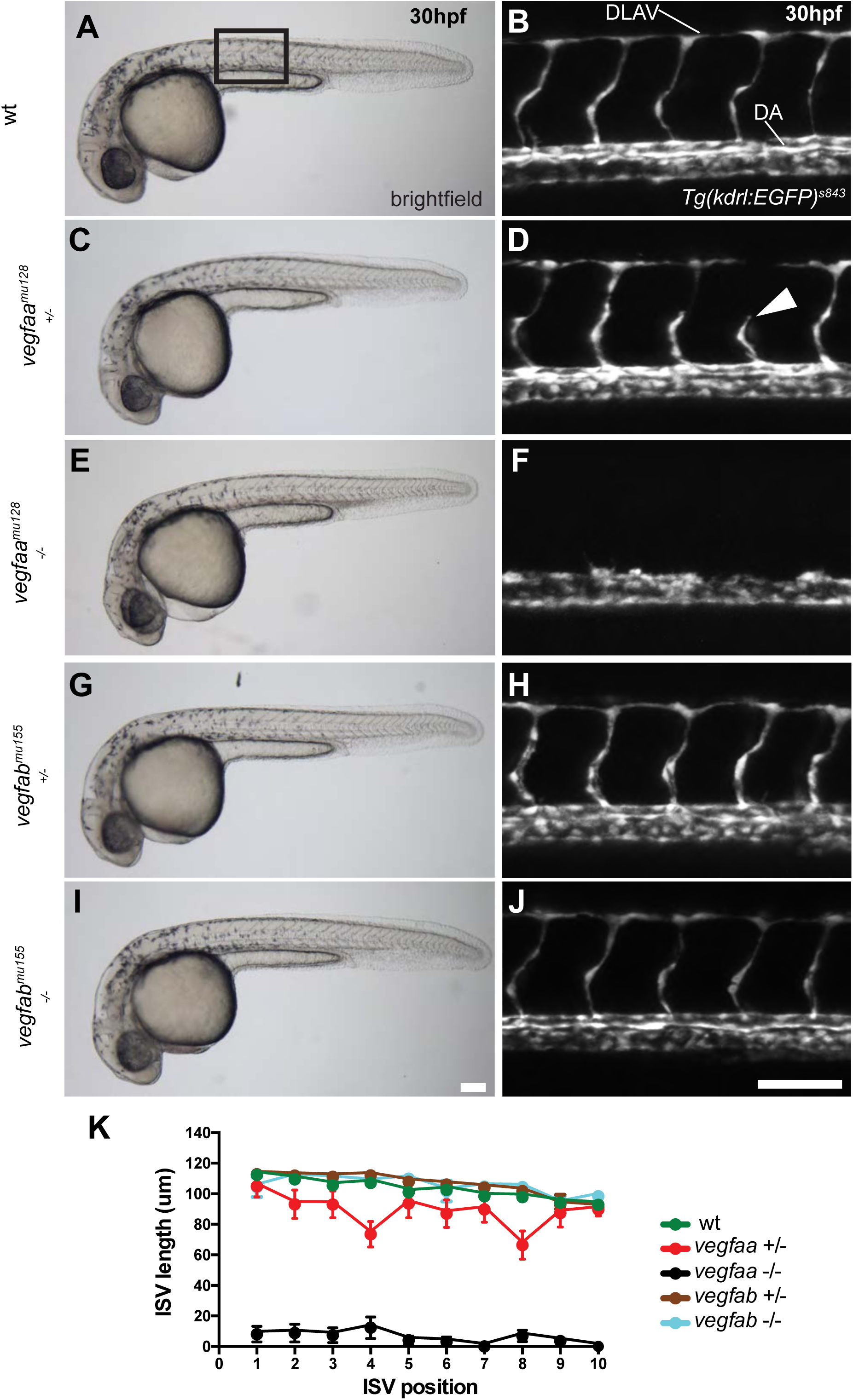
Analysis of overall morphology and ISV formation in vegfaa^mu128^ and vegfab^mu155^ mutants at 30 hpf. (A, C, E, G and I) Brightfield images of wild type (A), *vegfaa*^*mu128 +/-*^ (C), *vegfaa*^*mu128 -/-*^ (E) *vegfab*^*mu155 +/-*^ (G) and *vegfab*^*mu155 -/-*^ (I) embryos. (B, D, F, H and J) Maximum intensity projections of confocal z-stacks of the trunk vasculature of *Tg(kdrl:EGFP)*^*s843*^ wild type (B), *vegfaa*^*mu128 +/-*^ (D), *vegfaa*^*mu128 -/-*^ (F) *vegfab*^*mu155 +/-*^ (H) and *vegfab*^*mu155 -/-*^ (J) embryos. Lateral views, anterior to the left. Scale bar = 100 um. (K) Quantification of ISV length from ISV number 1 to number 10 from anterior to posterior of the trunk vasculature of wild type (green), *vegfaa*^*mu128 +/-*^ (red), *vegfaa*^*mu128 -/-*^ (black) *vegfab*^*mu155 +/-*^ (brown) and *vegfab*^*mu155 -/-*^ (blue) embryos; n = 8. Values are mean±s.d.

**Supplementary Figure S3.**
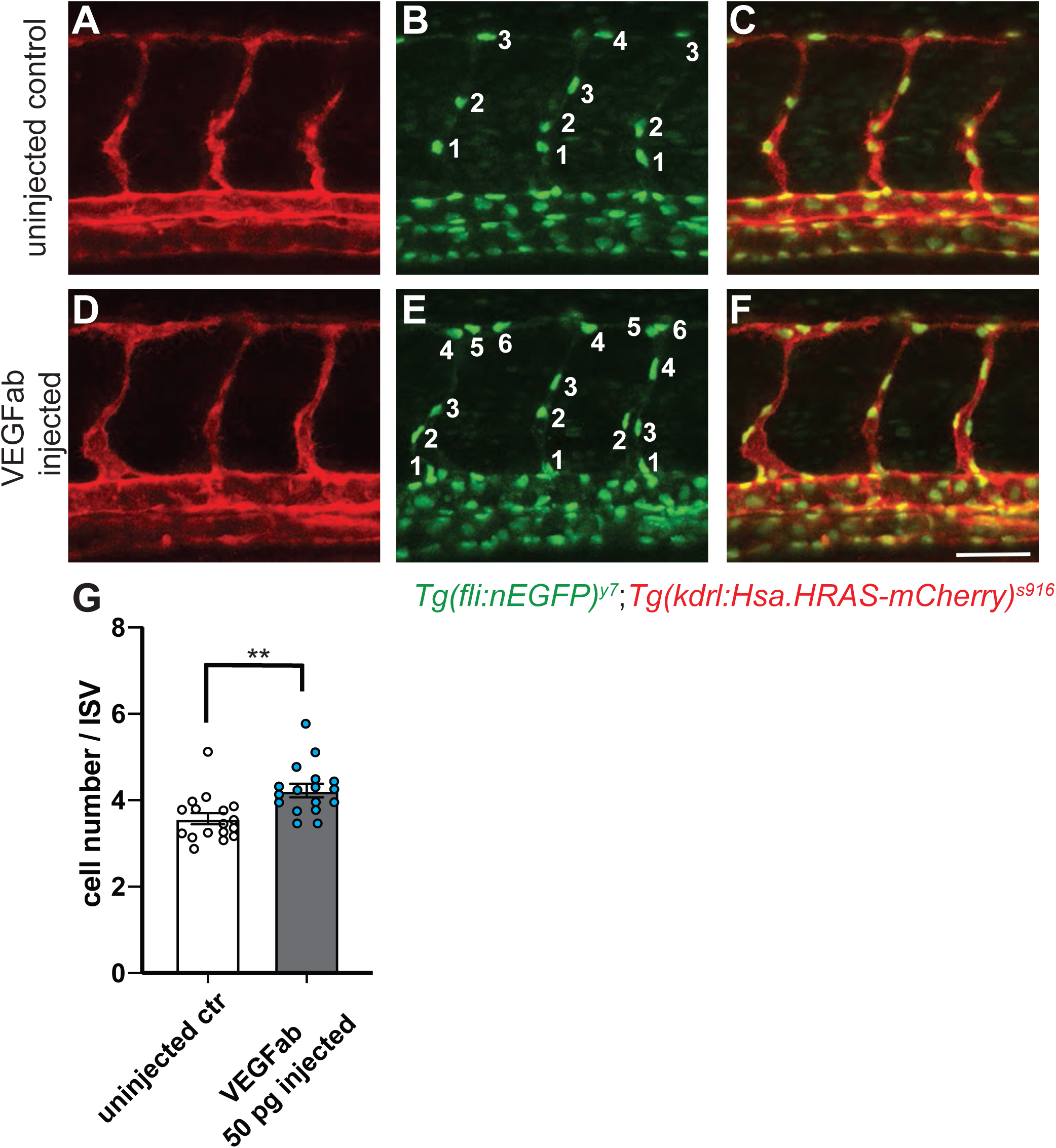
Overexpression of Vegfab leads to an increase in ISV cell numbers. (A-C) ISVs of uninjected control embryos at 32 hpf. Blood vessels labelled by *Tg(Hsa.HRAS-mCherry)*^*s916*^ in red (A). Endothelial cell nuclei labelled by *Tg(Fli:nEGFP)*^*y7*^ in green (B). Overlay of red and green channels (C). Side views, anterior to the left. (D-F) ISVs of embryos injected with 50pg of vegfab mRNA at 32 hpf. Blood vessels labelled by *Tg(Hsa.HRAS-mCherry)*^*s916*^ in red (D). Endothelial cell nuclei labelled by *Tg(Fli:nEGFP)*^*y7*^ in green (E). Overlay of red and green channels (F). Scale bar is 50 um. (G) Quantification of ISV cell numbers in control and vegfab mRNA injected embryos (n=17 embryos each).

**Supplementary Figure S4.**
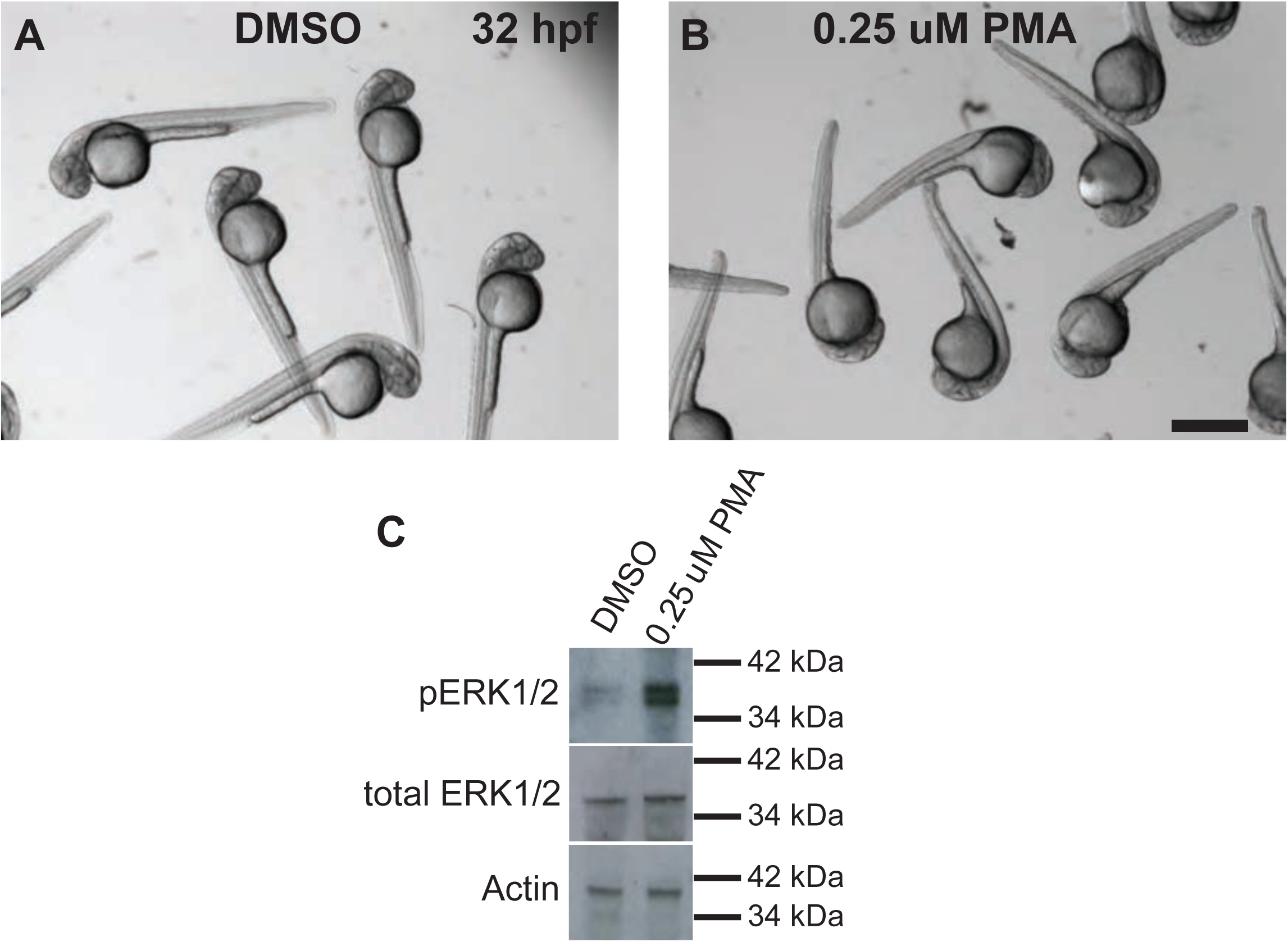
PMA treatment of zebrafish embryos leads to ERK phosphorylation. (A-B) Overview pictures of wild type embryos at 32 hpf treated with DMSO (A) or 0.25 uM PMA (B). Scale bar = 400 um. (C) Western blot analysis of pERK, total ERK and Actin in embryos treated with indicated concentrations of PMA or DMSO. Representative western blot of n=3 is shown.

**Supplementary Figure S5.**
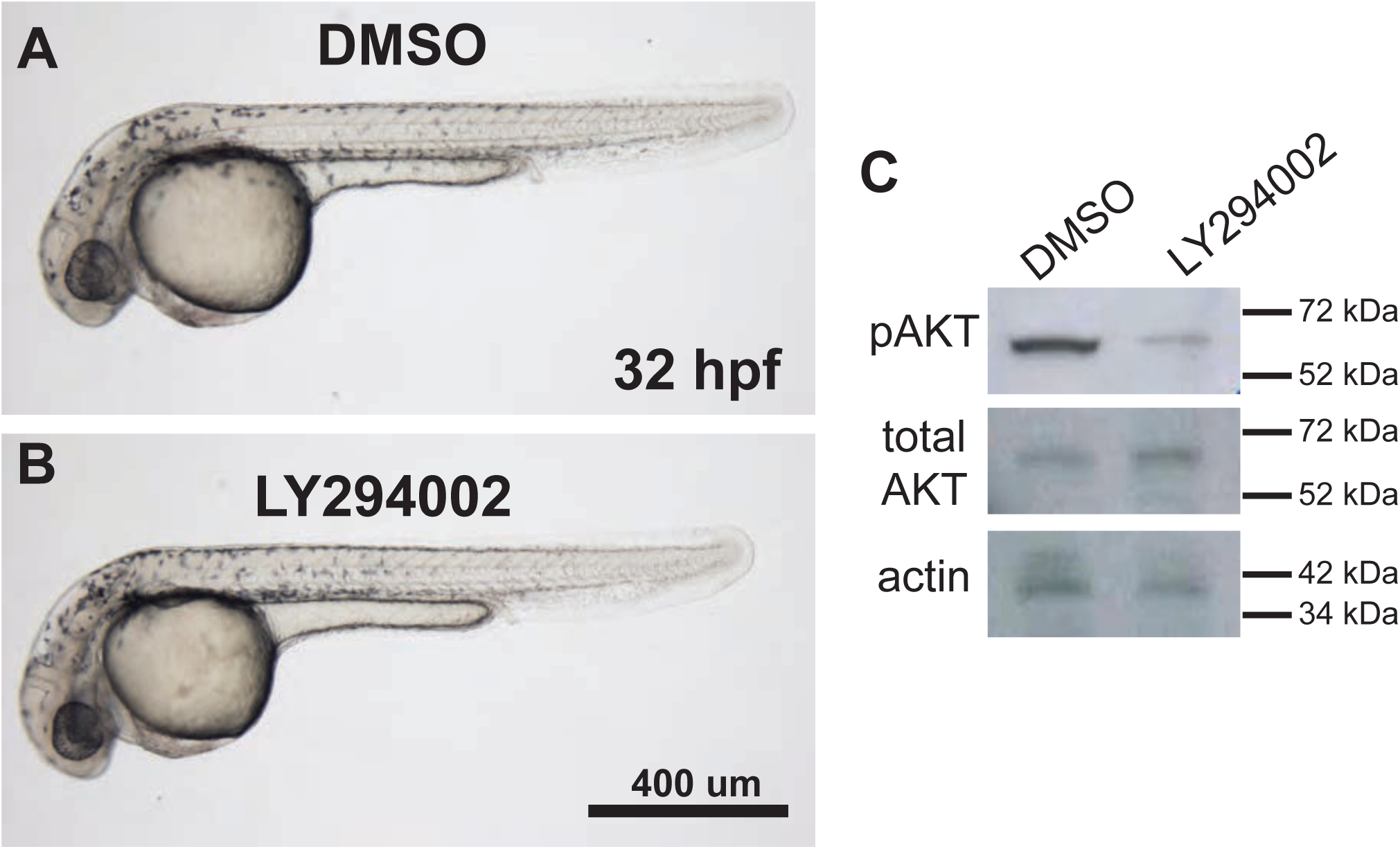
LY294002 treatment of zebrafish embryos leads to a reduction in Akt phosphorylation. (A-B) Brightfield images of single wild type embryos at 32 hpf treated with DMSO (A) or 10 uM LY294002 (B). Lateral views, anterior to the left. Scale bar = 400 um. (C) Western blot analysis of pAKT, total AKT and Actin in embryos treated with DMSO or 10 uM LY294002. Representative blot of n=3 is shown.

**Supplementary Figure S6.**
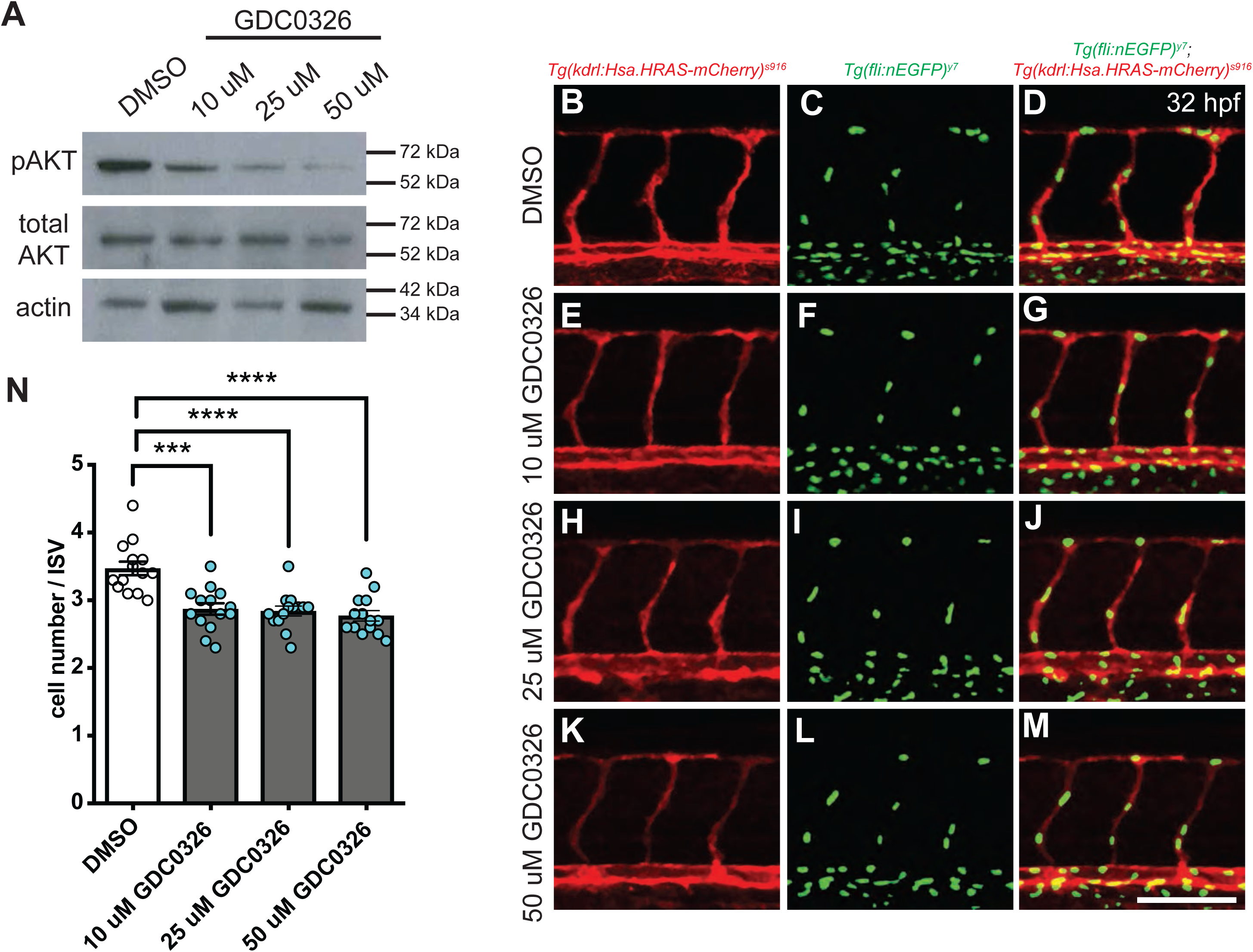
GDC-0326 mediated inhibition of PI3K p110 alpha isoform affects endothelial cell numbers. (A) Western blot analysis of pAKT, total AKT and Actin in embryos treated with DMSO or indicated concentrations of GDC-0326. Representative blot of n=3 is shown. (B-M) Maximum intensity projection of confocal z-stacks of the trunk vasculature of double transgenic *Tg(kdrl:Hsa.HRAS-mCherry)*^*s916*^; *Tg(fli1a:nEGFP)*^*y7*^ embryos treated with DMSO (B-D), 10 uM GDC-0326 (E-G), 25 uM GDC-0326 (H-J) or 50 uM GDC-0326 (K-M). Lateral views, anterior to the left. Scale bar = 100 um. (N) Quantification of cells per ISV in embryos treated with DMSO compared to embryos treated with 10 uM, 25 uM and 50 uM GDC-0326. Dots represent individual embryos (n = 14 embryos for each condition), black lines indicate the mean value ±s.d. *** p>0.0003; **** p<0.0001.

**Supplementary Figure S7.**
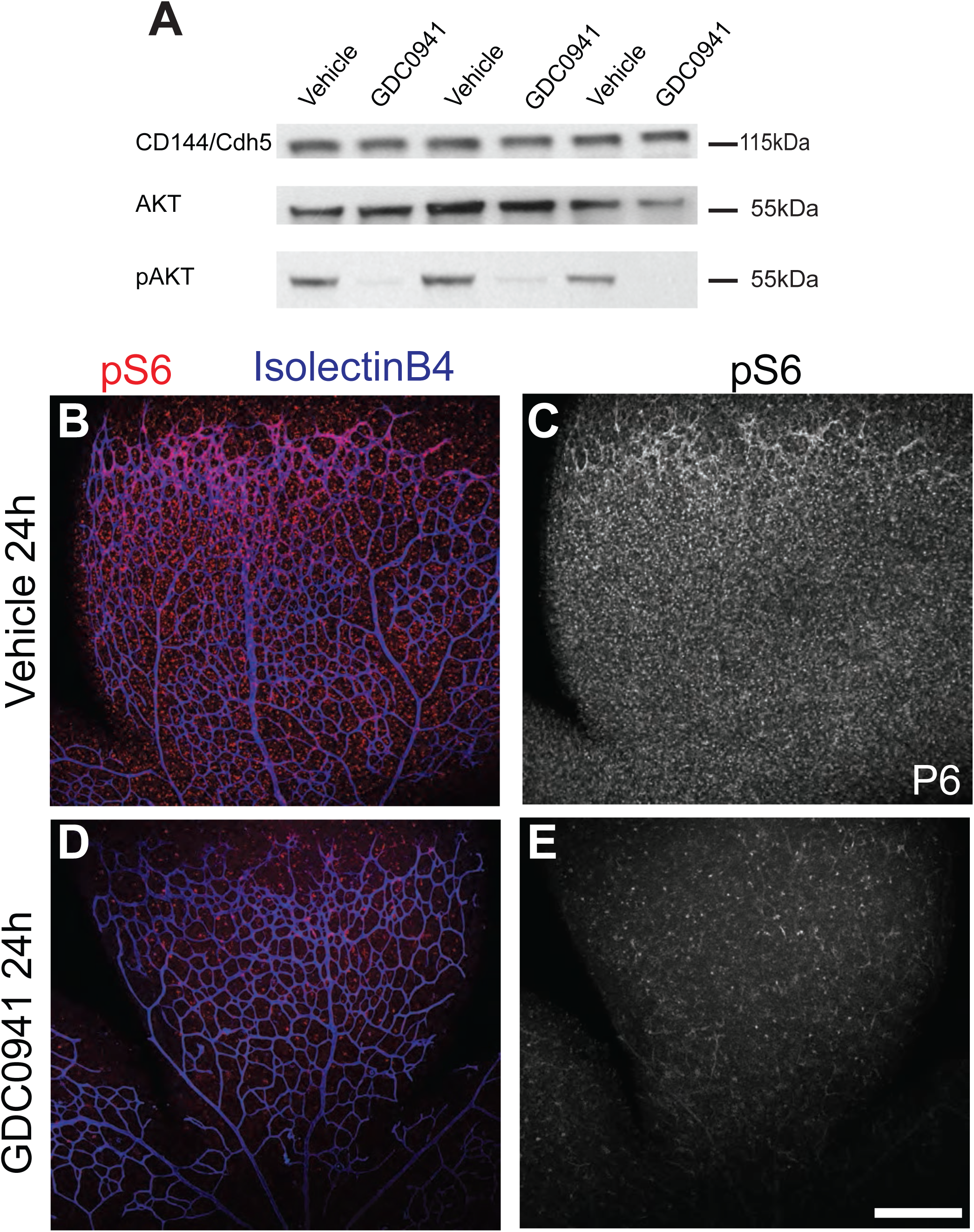
Treatment of mouse retinas with GDC-0941 leads to reduction in AKT phosphorylation and S6 phosphorylation. (A) Western blot analysis of pAKT, total AKT and CD144/Cdh5. Representative blot of n=3 is shown. (B-C) Immunohistochemistry on mouse retina. Staining for isolectinB4 (blue; blood vessels) and pS6 kinase (red) in vehicle injected control mice (B, C) and GDC-0941 injectected mice (D, E). Scale bar is 150 um.

**Supplementary Video S1. Development of hindbrain vasculature in wildtype zebrafish embryo**

Endothelial cells are marked by virtue of EGFP expression in *Tg(kdrl:EGFP)*^*s843*^ zebrafish. Dorsal view of head region, anterior to the left. Imaging starts at 28 hpf and is carried out for 18.5 hours.

**Supplementary Video S2. Development of hindbrain vasculature in *vegfaa***^***mu128***^ **zebrafish embryo**

Endothelial cells are marked by virtue of EGFP expression in *Tg(kdrl:EGFP)*^*s843*^ zebrafish. Dorsal view of head region, anterior to the left. Imaging starts at 28 hpf and is carried out for 18.5 hours.

**Supplementary Video S3. Development of hindbrain vasculature in *vegfab***^***mu155***^ **zebrafish embryo**

Endothelial cells are marked by virtue of EGFP expression in *Tg(kdrl:EGFP)*^*s843*^ zebrafish. Dorsal view of head region, anterior to the left. Imaging starts at 28 hpf and is carried out for 18.5 hours.

**Supplementary Video S4. Development of hindbrain vasculature in wildtype zebrafish embryo**

Endothelial cell membranes (red) are marked by virtue of mCherry expression, while endothelial cell nuclei express EGFP (green) in *Tg(kdrl:Hsa.HRAS-mCherry)*^*s916*^; *Tg(fli1:nEGFP)*^*y7*^ zebrafish. Dorsal view of head region, anterior to the left. Imaging starts at 28 hpf and is carried out for 14 hours.

**Supplementary Video S5. Development of hindbrain vasculature in *vegfaa***^***mu128***^ **zebrafish embryo**

Endothelial cell membranes (red) are marked by virtue of mCherry expression, while endothelial cell nuclei express EGFP (green) in *Tg(kdrl:Hsa.HRAS-mCherry)*^*s916*^; *Tg(fli1:nEGFP)*^*y7*^ zebrafish. Dorsal view of head region, anterior to the left. Imaging starts at 28 hpf and is carried out for 14 hours.

**Supplementary Video S6. Development of hindbrain vasculature in *vegfab***^***mu155***^ **zebrafish embryo**

Endothelial cell membranes (red) are marked by virtue of mCherry expression, while endothelial cell nuclei express EGFP (green) in *Tg(kdrl:Hsa.HRAS-mCherry)*^*s916*^; *Tg(fli1:nEGFP)*^*y7*^ zebrafish. Dorsal view of head region, anterior to the left. Imaging starts at 28 hpf and is carried out for 14.5 hours.

**Supplementary Video S7. Endothelial cell proliferation and migration in individual intersegmental blood vessel sprout in wildtype zebrafish embryo**

Endothelial cell membranes (red) are marked by virtue of mCherry expression, while endothelial cell nuclei express EGFP (green) in *Tg(kdrl:Hsa.HRAS-mCherry)*^*s916*^; *Tg(fli1:nEGFP)*^*y7*^ zebrafish. Side view of trunk region, anterior to the left. Imaging starts at 22 hpf and is carried out for 6 hours. Individual endothelial cells are marked by arrowheads and numbers. Decimal numbers indicate progeny after division.

**Supplementary Video S8. Endothelial cell proliferation and migration in individual intersegmental blood vessel sprout in *vegfaa***^***mu128***^ **heterozygous zebrafish embryo**

Endothelial cell membranes (red) are marked by virtue of mCherry expression, while endothelial cell nuclei express EGFP (green) in *Tg(kdrl:Hsa.HRAS-mCherry)*^*s916*^; *Tg(fli1:nEGFP)*^*y7*^ zebrafish. This example shows intersegmental vessel with normal endothelial cell behaviors. Side view of trunk region, anterior to the left. Imaging starts at 22 hpf and is carried out for 6 hours. Individual endothelial cells are marked by arrowheads and numbers. Decimal numbers indicate progeny after division.

**Supplementary Video S9. Endothelial cell proliferation and migration in individual intersegmental blood vessel sprout in *vegfaa***^***mu128***^ **heterozygous zebrafish embryo**

Endothelial cell membranes (red) are marked by virtue of mCherry expression, while endothelial cell nuclei express EGFP (green) in *Tg(kdrl:Hsa.HRAS-mCherry)*^*s916*^; *Tg(fli1:nEGFP)*^*y7*^ zebrafish. This example shows a stalled intersegmental vessel. Side view of trunk region, anterior to the left. Imaging starts at 22 hpf and is carried out for 6 hours. Individual endothelial cells are marked by arrowheads and numbers. Decimal numbers indicate progeny after division.

**Supplementary Video S10. Endothelial cell proliferation and migration in individual intersegmental blood vessel sprout in *vegfab***^***mu155***^ **mutant zebrafish embryo**

Endothelial cell membranes (red) are marked by virtue of mCherry expression, while endothelial cell nuclei express EGFP (green) in *Tg(kdrl:Hsa.HRAS-mCherry)*^*s916*^; *Tg(fli1:nEGFP)*^*y7*^ zebrafish Side view of trunk region, anterior to the left. Imaging starts at 22 hpf and is carried out for 6 hours. Individual endothelial cells are marked by arrowheads and numbers. Decimal numbers indicate progeny after division.

**Supplementary Video S11. Endothelial cell proliferation and migration in intersegmental blood vessel sprouts in wildtype zebrafish embryo**

Endothelial cell membranes (red) are marked by virtue of mCherry expression, while endothelial cell nuclei express EGFP (green) in *Tg(kdrl:Hsa.HRAS-mCherry)*^*s916*^; *Tg(fli1:nEGFP)*^*y7*^ zebrafish. Side view of trunk region, anterior to the left. Imaging starts at 22 hpf and is carried out for 10 hours. Individual endothelial cells are marked by arrowheads and numbers. Decimal numbers indicate progeny after division.

**Supplementary Video S12. Endothelial cell proliferation and migration in intersegmental blood vessel sprouts in *vegfaa***^***mu128***^ **heterozygous embryo**

Endothelial cell membranes (red) are marked by virtue of mCherry expression, while endothelial cell nuclei express EGFP (green) in *Tg(kdrl:Hsa.HRAS-mCherry)*^*s916*^; *Tg(fli1:nEGFP)*^*y7*^ zebrafish. Side view of trunk region, anterior to the left. Imaging starts at 22 hpf and is carried out for 10.5 hours. Individual endothelial cells are marked by arrowheads and numbers. Decimal numbers indicate progeny after division.

**Supplementary Video S13. Endothelial cell proliferation and migration in intersegmental blood vessel sprouts in *vegfab***^***mu155***^ **mutant embryo**

Endothelial cell membranes (red) are marked by virtue of mCherry expression, while endothelial cell nuclei express EGFP (green) in *Tg(kdrl:Hsa.HRAS-mCherry)*^*s916*^; *Tg(fli1:nEGFP)*^*y7*^ zebrafish. Side view of trunk region, anterior to the left. Imaging starts at 22 hpf and is carried out for 10.5 hours. Individual endothelial cells are marked by arrowheads and numbers. Decimal numbers indicate progeny after division.

**Supplementary Video S14. Endothelial cell proliferation and migration in individual intersegmental blood vessel sprout in DMSO treated zebrafish embryo**

Endothelial cell membranes (red) are marked by virtue of mCherry expression, while endothelial cell nuclei express EGFP (green) in *Tg(kdrl:Hsa.HRAS-mCherry)*^*s916*^; *Tg(fli1:nEGFP)*^*y7*^ zebrafish Side view of trunk region, anterior to the left. Imaging starts at 22 hpf and is carried out for 6 hours. Individual endothelial cells are marked by arrowheads and numbers. Decimal numbers indicate progeny after division.

**Supplementary Video S15. Endothelial cell proliferation and migration in individual intersegmental blood vessel sprout in LY294002 treated zebrafish embryo**

Endothelial cell membranes (red) are marked by virtue of mCherry expression, while endothelial cell nuclei express EGFP (green) in *Tg(kdrl:Hsa.HRAS-mCherry)*^*s916*^; *Tg(fli1:nEGFP)*^*y7*^ zebrafish Side view of trunk region, anterior to the left. Imaging starts at 22 hpf and is carried out for 6 hours. Individual endothelial cells are marked by arrowheads and numbers. Decimal numbers indicate progeny after division.

**Supplementary Video S16. Endothelial cell proliferation and migration in intersegmental blood vessel sprouts in DMSO treated zebrafish embryo**

Endothelial cell membranes (red) are marked by virtue of mCherry expression, while endothelial cell nuclei express EGFP (green) in *Tg(kdrl:Hsa.HRAS-mCherry)*^*s916*^; *Tg(fli1:nEGFP)*^*y7*^ zebrafish. Side view of trunk region, anterior to the left. Imaging starts at 22 hpf and is carried out for 10 hours. Individual endothelial cells are marked by arrowheads and numbers. Decimal numbers indicate progeny after division.

**Supplementary Video S17. Endothelial cell proliferation and migration in intersegmental blood vessel sprouts in LY294002 treated zebrafish embryo** Endothelial cell membranes (red) are marked by virtue of mCherry expression, while endothelial cell nuclei express EGFP (green) in *Tg(kdrl:Hsa.HRAS-mCherry)*^*s916*^; *Tg(fli1:nEGFP)*^*y7*^ zebrafish. Side view of trunk region, anterior to the left. Imaging starts at 22 hpf and is carried out for 10 hours. Individual endothelial cells are marked by arrowheads and numbers. Decimal numbers indicate progeny after division.

## REFERENCES

Adams, R.H., and Alitalo, K. (2007). Molecular regulation of angiogenesis and lymphangiogenesis. Nat Rev Mol Cell Biol 8, 464–478.

Alvarez-Aznar, A., Muhl, L., and Gaengel, K. (2017). VEGF Receptor Tyrosine Kinases: Key Regulators of Vascular Function. Curr Top Dev Biol 123, 433–482.

Angulo-Urarte, A., Casado, P., Castillo, S.D., Kobialka, P., Kotini, M.P., Figueiredo, A.M., Castel, P., Rajeeve, V., Mila-Guasch, M., Millan, J., et al. (2018). Endothelial cell rearrangements during vascular patterning require PI3- kinase-mediated inhibition of actomyosin contractility. Nature communications 9, 4826.

Bahary, N., Goishi, K., Stuckenholz, C., Weber, G., Leblanc, J., Schafer, C.A., Berman, S.S., Klagsbrun, M., and Zon, L.I. (2007). Duplicate VegfA genes and orthologues of the KDR receptor tyrosine kinase family mediate vascular development in the zebrafish. Blood 110, 3627–3636.

Bowler, E., and Oltean, S. (2019). Alternative Splicing in Angiogenesis. Int J Mol Sci 20.

Brash, J.T., Denti, L., Ruhrberg, C., and Bucher, F. (2019). VEGF188 promotes corneal reinnervation after injury. JCI Insight 4.

Bussmann, J., Wolfe, S.A., and Siekmann, A.F. (2011). Arterial-venous network formation during brain vascularization involves hemodynamic regulation of chemokine signaling. Development 138, 1717–1726.

Carmeliet, P., Ferreira, V., Breier, G., Pollefeyt, S., Kieckens, L., Gertsenstein, M., Fahrig, M., Vandenhoeck, A., Harpal, K., Eberhardt, C., et al. (1996). Abnormal blood vessel development and lethality in embryos lacking a single VEGF allele. Nature 380, 435–439.

Carmeliet, P., Ng, Y.S., Nuyens, D., Theilmeier, G., Brusselmans, K., Cornelissen, I., Ehler, E., Kakkar, V.V., Stalmans, I., Mattot, V., et al. (1999). Impaired myocardial angiogenesis and ischemic cardiomyopathy in mice lacking the vascular endothelial growth factor isoforms VEGF164 and VEGF188. Nat Med 5, 495–502.

Castel, P., Carmona, F.J., Grego-Bessa, J., Berger, M.F., Viale, A., Anderson, K.V., Bague, S., Scaltriti, M., Antonescu, C.R., Baselga, E., et al. (2016). Somatic PIK3CA mutations as a driver of sporadic venous malformations. Science translational medicine 8, 332ra342.

Castillo, S.D., Baselga, E., and Graupera, M. (2019). PIK3CA mutations in vascular malformations. Curr Opin Hematol 26, 170–178.

Castillo, S.D., Tzouanacou, E., Zaw-Thin, M., Berenjeno, I.M., Parker, V.E., Chivite, I., Mila-Guasch, M., Pearce, W., Solomon, I., Angulo-Urarte, A., et al. (2016). Somatic activating mutations in Pik3ca cause sporadic venous malformations in mice and humans. Science translational medicine 8, 332ra343.

Chen, T.T., Luque, A., Lee, S., Anderson, S.M., Segura, T., and Iruela-Arispe, M.L. (2010). Anchorage of VEGF to the extracellular matrix conveys differential signaling responses to endothelial cells. J Cell Biol 188, 595–609.

Cheng, S.Y., Nagane, M., Huang, H.S., and Cavenee, W.K. (1997). Intracerebral tumor-associated hemorrhage caused by overexpression of the vascular endothelial growth factor isoforms VEGF121 and VEGF165 but not VEGF189. Proc Natl Acad Sci U S A 94, 12081–12087.

Childs, S., Chen, J.N., Garrity, D.M., and Fishman, M.C. (2002). Patterning of angiogenesis in the zebrafish embryo. Development 129, 973–982.

Chung, J., Grammer, T.C., Lemon, K.P., Kazlauskas, A., and Blenis, J. (1994). PDGF- and insulin-dependent pp70S6k activation mediated by phosphatidylinositol-3-OH kinase. Nature 370, 71–75.

Claesson-Welsh, L. (2016). VEGF receptor signal transduction - A brief update. Vascul Pharmacol 86, 14-17.

Costa, G., Harrington, K.I., Lovegrove, H.E., Page, D.J., Chakravartula, S., Bentley, K., and Herbert, S.P. (2016). Asymmetric division coordinates collective cell migration in angiogenesis. Nature cell biology 18, 1292–1301.

Davies, S.P., Reddy, H., Caivano, M., and Cohen, P. (2000). Specificity and mechanism of action of some commonly used protein kinase inhibitors. Biochem J 351, 95–105.

Dayanir, V., Meyer, R.D., Lashkari, K., and Rahimi, N. (2001). Identification of tyrosine residues in vascular endothelial growth factor receptor-2/FLK-1 involved in activation of phosphatidylinositol 3-kinase and cell proliferation. J Biol Chem 276, 17686–17692.

Dejana, E., Hirschi, K.K., and Simons, M. (2017). The molecular basis of endothelial cell plasticity. Nature communications 8, 14361.

Delcombel, R., Janssen, L., Vassy, R., Gammons, M., Haddad, O., Richard, B., Letourneur, D., Bates, D., Hendricks, C., Waltenberger, J., et al. (2013). New prospects in the roles of the C-terminal domains of VEGF-A and their cooperation for ligand binding, cellular signaling and vessels formation. Angiogenesis 16, 353–371.

Fearnley, G.W., Smith, G.A., Abdul-Zani, I., Yuldasheva, N., Mughal, N.A., Homer-Vanniasinkam, S., Kearney, M.T., Zachary, I.C., Tomlinson, D.C., Harrison, M.A., et al. (2016). VEGF-A isoforms program differential VEGFR2 signal transduction, trafficking and proteolysis. Biology open 5, 571–583.

Ferrara, N., Carver-Moore, K., Chen, H., Dowd, M., Lu, L., O’Shea, K.S., Powell-Braxton, L., Hillan, K.J., and Moore, M.W. (1996). Heterozygous embryonic lethality induced by targeted inactivation of the VEGF gene. Nature 380, 439–442.

Fish, J.E., Cantu Gutierrez, M., Dang, L.T., Khyzha, N., Chen, Z., Veitch, S., Cheng, H.S., Khor, M., Antounians, L., Njock, M.S., et al. (2017). Dynamic regulation of VEGF-inducible genes by an ERK/ERG/p300 transcriptional network. Development 144, 2428–2444.

Fujita, M., Cha, Y.R., Pham, V.N., Sakurai, A., Roman, B.L., Gutkind, J.S., and Weinstein, B.M. (2011). Assembly and patterning of the vascular network of the vertebrate hindbrain. Development 138, 1705–1715.

Gerber, H.P., McMurtrey, A., Kowalski, J., Yan, M., Keyt, B.A., Dixit, V., and Ferrara, N. (1998). Vascular endothelial growth factor regulates endothelial cell survival through the phosphatidylinositol 3’-kinase/Akt signal transduction pathway. Requirement for Flk-1/KDR activation. J Biol Chem 273, 30336–30343.

Gimbrone, M.A., Jr., and Garcia-Cardena, G. (2016). Endothelial Cell Dysfunction and the Pathobiology of Atherosclerosis. Circ Res 118, 620–636.

Gong, B., Liang, D., Chew, T.G., and Ge, R. (2004). Characterization of the zebrafish vascular endothelial growth factor A gene: comparison with vegf-A genes in mammals and Fugu. Biochimica et biophysica acta 1676, 33-40.

Graupera, M., Guillermet-Guibert, J., Foukas, L.C., Phng, L.K., Cain, R.J., Salpekar, A., Pearce, W., Meek, S., Millan, J., Cutillas, P.R., et al. (2008). Angiogenesis selectively requires the p110alpha isoform of PI3K to control endothelial cell migration. Nature 453, 662–666.

Graupera, M., and Potente, M. (2013). Regulation of angiogenesis by PI3K signaling networks. Exp Cell Res 319, 1348–1355.

Guo, P., Xu, L., Pan, S., Brekken, R.A., Yang, S.T., Whitaker, G.B., Nagane, M., Thorpe, P.E., Rosenbaum, J.S., Su Huang, H.J., et al. (2001). Vascular endothelial growth factor isoforms display distinct activities in promoting tumor angiogenesis at different anatomic sites. Cancer Res 61, 8569–8577.

Hasan, S.S., Tsaryk, R., Lange, M., Wisniewski, L., Moore, J.C., Lawson, N.D., Wojciechowska, K., Schnittler, H., and Siekmann, A.F. (2017). Endothelial Notch signalling limits angiogenesis via control of artery formation. Nature cell biology 19, 928–940.

Heffron, T.P., Heald, R.A., Ndubaku, C., Wei, B., Augistin, M., Do, S., Edgar, K., Eigenbrot, C., Friedman, L., Gancia, E., et al. (2016). The Rational Design of Selective Benzoxazepin Inhibitors of the alpha-Isoform of Phosphoinositide 3-Kinase Culminating in the Identification of (S)-2-((2-(1-Isopropyl-1H-1,2,4-triazol-5-yl)-5,6- dihydrobenzo[f]imidazo[1,2-d][1, 4]oxazepin-9-yl)oxy)propanamide (GDC-0326). J Med Chem 59, 985–1002.

Herve, M.A., Buteau-Lozano, H., Mourah, S., Calvo, F., and Perrot-Applanat, M. (2005). VEGF189 stimulates endothelial cells proliferation and migration in vitro and up-regulates the expression of Flk-1/KDR mRNA. Exp Cell Res 309, 24–31.

Hogan, B.M., Bos, F.L., Bussmann, J., Witte, M., Chi, N.C., Duckers, H.J., and Schulte-Merker, S. (2009). ccbe1 is required for embryonic lymphangiogenesis and venous sprouting. Nature Genetics 41, 396–398.

Hogan, B.M., and Schulte-Merker, S. (2017). How to Plumb a Pisces: Understanding Vascular Development and Disease Using Zebrafish Embryos. Dev Cell 42, 567–583.

Isogai, S., Lawson, N.D., Torrealday, S., Horiguchi, M., and Weinstein, B.M. (2003). Angiogenic network formation in the developing vertebrate trunk. Development 130, 5281–5290.

Janetopoulos, C., Borleis, J., Vazquez, F., Iijima, M., and Devreotes, P. (2005). Temporal and spatial regulation of phosphoinositide signaling mediates cytokinesis. Dev Cell 8, 467–477.

Jin, D., Zhu, D., Fang, Y., Chen, Y., Yu, G., Pan, W., Liu, D., Li, F., and Zhong, T.P. (2017). Vegfa signaling regulates diverse artery/vein formation in vertebrate vasculatures. Journal of genetics and genomics = Yi chuan xue bao 44, 483–492.

Jin, S.W., Beisl, D., Mitchell, T., Chen, J.N., and Stainier, D.Y.R. (2005). Cellular and molecular analyses of vascular tube and lumen formation in zebrafish. Development 132, 5199–5209.

Kazemi, M., Carrer, A., Moimas, S., Zandona, L., Bussani, R., Casagranda, B., Palmisano, S., Prelazzi, P., Giacca, M., Zentilin, L., et al. (2016). VEGF121 and VEGF165 differentially promote vessel maturation and tumor growth in mice and humans. Cancer Gene Ther 23, 125–132.

Koch, S., and Claesson-Welsh, L. (2012). Signal transduction by vascular endothelial growth factor receptors. Cold Spring Harbor perspectives in medicine 2, a006502.

Kochhan, E., and Siekmann, A.F. (2013). Zebrafish as a Model to Study Chemokine Function. In Chemokines: Methods and Protocols, E.A. Cardona, and E.E. Ubogu, eds. (Totowa, NJ: Humana Press), pp. 145–159.

Kreis, N.N., Louwen, F., and Yuan, J. (2019). The Multifaceted p21 (Cip1/Waf1/CDKN1A) in Cell Differentiation, Migration and Cancer Therapy. Cancers (Basel) 11.

Lawson, N.D., Scheer, N., Pham, V.N., Kim, C.H., Chitnis, A.B., Campos-Ortega, J.A., and Weinstein, B.M. (2001). Notch signaling is required for arterial-venous differentiation during embryonic vascular development. Development 128, 3675–3683.

Lawson, N.D., Vogel, A.M., and Weinstein, B.M. (2002). sonic hedgehog and vascular endothelial growth factor act upstream of the Notch pathway during arterial endothelial differentiation. Dev Cell 3, 127–136.

Liang, D., Xu, X., Chin, A.J., Balasubramaniyan, N.V., Teo, M.A., Lam, T.J., Weinberg, E.S., and Ge, R. (1998). Cloning and characterization of vascular endothelial growth factor (VEGF) from zebrafish, Danio rerio. Biochimica et biophysica acta 1397, 14–20.

Mavria, G., Vercoulen, Y., Yeo, M., Paterson, H., Karasarides, M., Marais, R., Bird, D., and Marshall, C.J. (2006). ERK-MAPK signaling opposes Rho-kinase to promote endothelial cell survival and sprouting during angiogenesis. Cancer Cell 9, 33–44.

Meadows, K.N., Bryant, P., and Pumiglia, K. (2001). Vascular endothelial growth factor induction of the angiogenic phenotype requires Ras activation. J Biol Chem 276, 49289–49298.

Mitchell, P., Liew, G., Gopinath, B., and Wong, T.Y. (2018). Age-related macular degeneration. Lancet 392, 1147–1159.

Nakatsu, M.N., Sainson, R.C., Perez-del-Pulgar, S., Aoto, J.N., Aitkenhead, M., Taylor, K.L., Carpenter, P.M., and Hughes, C.C. (2003). VEGF(121) and VEGF(165) regulate blood vessel diameter through vascular endothelial growth factor receptor 2 in an in vitro angiogenesis model. Lab Invest 83, 1873–1885.

Nicoli, S., Knyphausen, C.P., Zhu, L.J., Lakshmanan, A., and Lawson, N.D. (2012). miR-221 is required for endothelial tip cell behaviors during vascular development. Dev Cell 22, 418–429.

Ola, R., Dubrac, A., Han, J., Zhang, F., Fang, J.S., Larrivee, B., Lee, M., Urarte, A.A., Kraehling, J.R., Genet, G., et al. (2016). PI3 kinase inhibition improves vascular malformations in mouse models of hereditary haemorrhagic telangiectasia. Nature communications 7, 13650.

Overton, K.W., Spencer, S.L., Noderer, W.L., Meyer, T., and Wang, C.L. (2014). Basal p21 controls population heterogeneity in cycling and quiescent cell cycle states. Proc Natl Acad Sci U S A 111, E4386–4393.

Pack, L.R., Daigh, L.H., and Meyer, T. (2019). Putting the brakes on the cell cycle: mechanisms of cellular growth arrest. Curr Opin Cell Biol 60, 106–113.

Pages, G., Guerin, S., Grall, D., Bonino, F., Smith, A., Anjuere, F., Auberger, P., and Pouyssegur, J. (1999). Defective thymocyte maturation in p44 MAP kinase (Erk 1) knockout mice. Science 286, 1374–1377.

Park, J.E., Keller, G.A., and Ferrara, N. (1993). The vascular endothelial growth factor (VEGF) isoforms: differential deposition into the subepithelial extracellular matrix and bioactivity of extracellular matrix-bound VEGF. Mol Biol Cell 4, 1317–1326.

Peach, C.J., Mignone, V.W., Arruda, M.A., Alcobia, D.C., Hill, S.J., Kilpatrick, L.E., and Woolard, J. (2018). Molecular Pharmacology of VEGF-A Isoforms: Binding and Signalling at VEGFR2. Int J Mol Sci 19.

Pontes-Quero, S., Fernandez-Chacon, M., Luo, W., Lunella, F.F., Casquero-Garcia, V., Garcia-Gonzalez, I., Hermoso, A., Rocha, S.F., Bansal, M., and Benedito, R. (2019). High mitogenic stimulation arrests angiogenesis. Nature communications 10, 2016.

Potente, M., Gerhardt, H., and Carmeliet, P. (2011). Basic and therapeutic aspects of angiogenesis. Cell 146, 873– 887.

Qian, X., Hulit, J., Suyama, K., Eugenin, E.A., Belbin, T.J., Loudig, O., Smirnova, T., Zhou, Z.N., Segall, J., Locker, J., et al. (2013). p21CIP1 mediates reciprocal switching between proliferation and invasion during metastasis. Oncogene 32, 2292–2303 e2297.

Ricard, N., Scott, R.P., Booth, C.J., Velazquez, H., Cilfone, N.A., Baylon, J.L., Gulcher, J.R., Quaggin, S.E., Chittenden, T.W., and Simons, M. (2019). Endothelial ERK1/2 signaling maintains integrity of the quiescent endothelium. J Exp Med 216, 1874–1890.

Roman, B.L., Pham, V.N., Lawson, N.D., Kulik, M., Childs, S., Lekven, A.C., Garrity, D.M., Moon, R.T., Fishman, M.C., Lechleider, R.J., et al. (2002). Disruption of acvrl1 increases endothelial cell number in zebrafish cranial vessels. Development 129, 3009–3019.

Rossi, A., Gauvrit, S., Marass, M., Pan, L., Moens, C.B., and Stainier, D.Y.R. (2016). Regulation of Vegf signaling by natural and synthetic ligands. Blood 128, 2359–2366.

Ruhrberg, C., Gerhardt, H., Golding, M., Watson, R., Ioannidou, S., Fujisawa, H., Betsholtz, C., and Shima, D.T. (2002). Spatially restricted patterning cues provided by heparin-binding VEGF-A control blood vessel branching morphogenesis. Genes Dev 16, 2684–2698.

Schuermann, A., Helker, C.S., and Herzog, W. (2014). Angiogenesis in zebrafish. Semin Cell Dev Biol 31, 106-114.

Shin, M., Beane, T.J., Quillien, A., Male, I., Zhu, L.J., and Lawson, N.D. (2016). Vegfa signals through ERK to promote angiogenesis, but not artery differentiation. Development 143, 3796–3805.

Shiying, W., Boyun, S., Jianye, Y., Wanjun, Z., Ping, T., Jiang, L., and Hongyi, H. (2017). The Different Effects of VEGFA121 and VEGFA165 on Regulating Angiogenesis Depend on Phosphorylation Sites of VEGFR2. Inflamm Bowel Dis 23, 603–616.

Sidi, S., Sanda, T., Kennedy, R.D., Hagen, A.T., Jette, C.A., Hoffmans, R., Pascual, J., Imamura, S., Kishi, S., Amatruda, J.F., et al. (2008). Chk1 suppresses a caspase-2 apoptotic response to DNA damage that bypasses p53, Bcl-2, and caspase-3. Cell 133, 864–877.

Siekmann, A.F., and Lawson, N.D. (2007). Notch signalling limits angiogenic cell behaviour in developing zebrafish arteries. Nature 445, 781–784.

Siekmann, A.F., Standley, C., Fogarty, K.E., Wolfe, S.A., and Lawson, N.D. (2009). Chemokine signaling guides regional patterning of the first embryonic artery. Gene Dev 23, 2272–2277.

Simons, M., and Eichmann, A. (2015). Molecular controls of arterial morphogenesis. Circ Res 116, 1712–1724.

Simons, M., Gordon, E., and Claesson-Welsh, L. (2016). Mechanisms and regulation of endothelial VEGF receptor signalling. Nat Rev Mol Cell Biol 17, 611–625.

Spencer, S.L., Cappell, S.D., Tsai, F.C., Overton, K.W., Wang, C.L., and Meyer, T. (2013). The proliferation-quiescence decision is controlled by a bifurcation in CDK2 activity at mitotic exit. Cell 155, 369–383.

Srinivasan, R., Zabuawala, T., Huang, H., Zhang, J., Gulati, P., Fernandez, S., Karlo, J.C., Landreth, G.E., Leone, G., and Ostrowski, M.C. (2009). Erk1 and Erk2 regulate endothelial cell proliferation and migration during mouse embryonic angiogenesis. PLoS One 4, e8283.

Stalmans, I., Ng, Y.S., Rohan, R., Fruttiger, M., Bouche, A., Yuce, A., Fujisawa, H., Hermans, B., Shani, M., Jansen, S., et al. (2002). Arteriolar and venular patterning in retinas of mice selectively expressing VEGF isoforms. J Clin Invest 109, 327–336.

Sugden, W.W., Meissner, R., Aegerter-Wilmsen, T., Tsaryk, R., Leonard, E.V., Bussmann, J., Hamm, M.J., Herzog, W., Jin, Y., Jakobsson, L., et al. (2017). Endoglin controls blood vessel diameter through endothelial cell shape changes in response to haemodynamic cues. Nature cell biology 19, 653–665.

Takahashi, T., Ueno, H., and Shibuya, M. (1999). VEGF activates protein kinase C-dependent, but Ras-independent Raf-MEK-MAP kinase pathway for DNA synthesis in primary endothelial cells. Oncogene 18, 2221–2230.

Tanaka, H., Yamashita, T., Asada, M., Mizutani, S., Yoshikawa, H., and Tohyama, M. (2002). Cytoplasmic p21(Cip1/WAF1) regulates neurite remodeling by inhibiting Rho-kinase activity. J Cell Biol 158, 321–329.

Taylor, J.S., Braasch, I., Frickey, T., Meyer, A., and Van de Peer, Y. (2003). Genome duplication, a trait shared by 22000 species of ray-finned fish. Genome research 13, 382–390.

Taylor, J.S., Van de Peer, Y., Braasch, I., and Meyer, A. (2001). Comparative genomics provides evidence for an ancient genome duplication event in fish. Philos Trans R Soc Lond B Biol Sci 356, 1661–1679.

Testini, C., Smith, R.O., Jin, Y., Martinsson, P., Sun, Y., Hedlund, M., Sainz-Jaspeado, M., Shibuya, M., Hellstrom, M., and Claesson-Welsh, L. (2019). Myc-dependent endothelial proliferation is controlled by phosphotyrosine 1212 in VEGF receptor-2. EMBO Rep 20, e47845.

Thisse, C., and Thisse, B. (2008). High-resolution in situ hybridization to whole-mount zebrafish embryos. Nat Protoc 3, 59–69.

Ulrich, F., Ma, L.H., Baker, R.G., and Torres-Vazquez, J. (2011). Neurovascular development in the embryonic zebrafish hindbrain. Dev Biol 357, 134–151.

Vempati, P., Popel, A.S., and Mac Gabhann, F. (2014). Extracellular regulation of VEGF: isoforms, proteolysis, and vascular patterning. Cytokine Growth Factor Rev 25, 1–19.

Viallard, C., and Larrivee, B. (2017). Tumor angiogenesis and vascular normalization: alternative therapeutic targets. Angiogenesis 20, 409–426.

Villefranc, J.A., Amigo, J., and Lawson, N.D. (2007). Gateway compatible vectors for analysis of gene function in the zebrafish. Dev Dyn 236, 3077–3087.

Vlahos, C.J., Matter, W.F., Hui, K.Y., and Brown, R.F. (1994). A specific inhibitor of phosphatidylinositol 3-kinase, 2-(4-morpholinyl)-8-phenyl-4H-1-benzopyran-4-one (LY294002). J Biol Chem 269, 5241–5248.

Westerfield, M. (1993). The zebrafish book : a guide for the laboratory use of zebrafish (Brachydanio rerio) (Eugene, OR: M. Westerfield).

Wong, T.Y., Cheung, C.M., Larsen, M., Sharma, S., and Simo, R. (2016). Diabetic retinopathy. Nat Rev Dis Primers 2, 16012.

Woolard, J., Bevan, H.S., Harper, S.J., and Bates, D.O. (2009). Molecular diversity of VEGF-A as a regulator of its biological activity. Microcirculation 16, 572–592.

Workman, P., Clarke, P.A., Raynaud, F.I., and van Montfort, R.L. (2010). Drugging the PI3 kinome: from chemical tools to drugs in the clinic. Cancer Res 70, 2146–2157.

Zhou, B.P., Liao, Y., Xia, W., Spohn, B., Lee, M.H., and Hung, M.C. (2001). Cytoplasmic localization of p21Cip1/WAF1 by Akt-induced phosphorylation in HER-2/neu-overexpressing cells. Nature cell biology 3, 245– 252.

